# Opto-CLIP reveals dynamic FMRP regulation of mRNAs upon CA1 neuronal activation

**DOI:** 10.1101/2024.08.13.607210

**Authors:** Ruth A. Singer, Veronika Rajchin, Kwanghoon Park, Nathaniel Heintz, Robert B. Darnell

**Author notes:** The Herbert Wertheim UF Scripps Institute for Biomedical Innovation & Technology, Jupiter, FL, USA.

## Abstract

Neuronal diversity and function are intricately linked to the dynamic regulation of RNA metabolism. Electrophysiologic studies of synaptic plasticity, models for learning and memory, are disrupted in Fragile X Syndrome (FXS). FXS is characterized by the loss of FMRP, an RNA-binding protein (RBP) known to suppress translation of specific neuronal RNAs. Synaptic plasticity in CA1 excitatory hippocampal neurons is protein-synthesis dependent, suggesting a role for FMRP in FXS-related synaptic deficits. To explore this model, we developed Opto-CLIP, integrating optogenetics with cell-type specific FMRP-CLIP and RiboTag in CA1 neurons, allowing investigation of activity-induced FMRP regulation. We tracked changes in FMRP binding and ribosome-associated RNA profiles 30 minutes after neuronal activation. Our findings reveal distinct temporal dynamics for FMRP transcript regulation in the cell body versus the synapse. In the cell body, FMRP binding to transcripts encoding nuclear functions is relieved, potentially allowing rapid transcriptional responses to neuronal activation. At the synapse, FMRP binding to transcripts encoding synaptic targets was relatively stable, with variability in translational control across target categories. These results offer fresh insights into the dynamic regulation of RNA by FMRP in response to neuronal activation and provide a foundation for future research into the mechanisms of RBP-mediated synaptic plasticity.

## INTRODUCTION

The uniformity of DNA across different cell types underscores the critical role of RNA transcription and processing in driving cellular diversity. Examining how RNA is spliced, polyadenylated, localized, and translated provides key insights into complex cellular functions. In neurons, RNA regulation has been of special interest to neuroscientists following the discovery that memory is protein synthesis dependent. The hippocampus, essential for new memory formation, serves as a model for studying the cellular basis of learning and memory.

Synaptic plasticity—the ability of synapses to strengthen or weaken over time in response to activity—is central to memory formation (Hughes 1958). In hippocampal CA1 pyramidal neurons, synaptic plasticity can be demonstrated by electrophysiologic induction of long-term potentiation (LTP) and long-term depression (LTD). Importantly, LTP induction in CA1 neurons is protein synthesis dependent and occurs locally within dendrites (Frey et al. 1989; Kang and Schuman 1996).

Dysregulation of synaptic plasticity has been implicated in neurodevelopmental disorders, including Fragile X Syndrome (FXS) (Bear et al. 2004; Huber et al. 2002). FXS, the most common inherited cause of intellectual disability and leading monogenic cause of autism, results from the silencing of the *Fmr1* gene and subsequent loss of its protein product, the RNA binding protein (RBP), FMRP (Hagerman et al. 2017; Ashley et al. 1993). Decades of research have demonstrated that FMRP is required for proper neurophysiological function. *Fmr1*-KO mice exhibit defects in cognition and learning/memory formation (Huber et al. 2002; Arbab et al. 2018; Boone et al. 2018; Talbot et al. 2018), dendritic spine morphology and dynamics (Comery et al. 1997), and synaptic plasticity (Lauterborn et al. 2007; Lee et al. 2011; Muddashetty et al. 2007), closely paralleling phenotypes observed in human FXS (Kazdoba et al. 2014). These findings have led to a model in which dysregulation of FMRP target RNAs disrupts the balance of glutamate-dependent excitation/ inhibition, underlying the development of intellectual disability in FXS (Bear et al. 2004; Huber et al. 2002, 2000). Thus, identifying the transcripts FMRP binds and regulates is fundamental to our understanding of synaptic plasticity and the pathophysiology of FXS (Darnell 2020).

Direct FMRP targets have been identified using Crosslinking Immunoprecipitation (CLIP) in whole mouse brain (Darnell et al. 2011; Korb et al. 2017), CA1 excitatory hippocampal neurons (Sawicka et al. 2019), and microdissected CA1 neurites and cell bodies (Hale et al. 2021). CLIP studies revealed that FMRP binds across the coding sequence of mRNAs, and biochemical evidence show that FMRP stalls ribosomes on these transcripts, indicating a direct role for FMRP in translational regulation (Darnell et al. 2011; Sawicka et al. 2019; Hale et al. 2021; Anadolu et al. 2023). Comparative analysis of FMRP binding and regulation revealed enrichment of transcripts related to Autism Spectrum Disorders (Darnell et al. 2011; Sawicka et al. 2019) and distinct regulation of synaptic and nuclear targets within the dendritic and cell body layers, respectively, of CA1 neurons (Hale et al. 2021). These studies illustrate how detailed analysis of RBP-RNA interactions provide valuable insights into mechanisms underlying intellectual disability in FXS. However, these studies have been conducted under steady state conditions, leaving the precise mechanisms by which FMRP globally regulates RNA in the dynamic context of synaptic plasticity incompletely understood.

Synaptic plasticity underlying learning and memory has historically been studied through physiological (French et al. 2001; Park et al. 2006), behavioral (Guzowski et al. 1999; Jaeger et al. 2018; Cavallaro et al. 2002), and chemical (Kim et al. 2010; Chen et al. 2017) approaches, followed by searches for genes with modified expression. While these studies identified activity-induced transcriptional changes, such as immediate-early genes, the observed gene expression changes varied widely across brain regions, times post activation, and stimulation paradigms (Cavallaro et al. 2002; Benito and Barco 2015). The use of optogenetics has revolutionized the field of neuroscience by enabling precise observation and control of cellular activities (Scanziani and Häusser 2009; Deisseroth et al. 2006).

Optogenetics offers several advantages for studying synaptic plasticity: it is rapid, reversible, cell-type specific, effective across short or long timescales, and it can be integrated with mouse models (Scanziani and Häusser 2009; Yizhar et al. 2011). The rapid timescale of optogenetics is especially valuable for studying FMRP, which is dephosphorylated within minutes of neuronal activation (Lee et al. 2011; Narayanan et al. 2007; Nalavadi et al. 2012). Leveraging optogenetics has the potential to uncover molecular mechanisms of protein synthesis dependent synaptic plasticity, providing insights into dynamic RNA and protein regulatory processes.

Here, we present the development and application of Opto-CLIP, a new platform for cell-type specific identification of RNAs regulated by FMRP in optogenetically activated excitatory CA1 hippocampal neurons. Our approach leverages Cre-dependent expression of channelrhodopsin (ChR2), GFP-tagged FMRP, and HA-tagged ribosomes specifically in CA1 neurons, enabling the study of RNA-protein interactions and ribosome-associated RNA profiles following optogenetic stimulation. By integrating data from Opto-FMRP-CLIP and Opto-RiboTag techniques, we developed a comprehensive model of activity-induced FMRP-mediated RNA regulation. These findings provide new insight into the dynamic regulation of RNA by FMRP in response to neuronal activation and provide a foundation for future research into the molecular mechanisms underlying learning, memory, and neurodevelopmental disorders.

## RESULTS

### Optogenetic activation of excitatory CA1 neurons of the mouse hippocampus

To enable molecular analyses of FMRP-bound transcripts specifically in excitatory hippocampal neurons, we crossed *Camk2a*-Cre mice (Tsien et al. 1996) to either conditionally tagged (cTag)-FMRP mice (Sawicka et al. 2019; Van Driesche et al. 2019) or *Rpl22*-HA (RiboTag) mice (Sanz et al. 2009). This strategy enables cell-type specific expression of either GFP tagged FMRP (*Fmr1*-cTag) or HA tagged ribosomes (RiboTag). Previous studies found nearly identical CLIP results with GFP-tagged FMRP and endogenous FMRP, supporting the use of the *Fmr1*-cTag mouse for CLIP studies (Van Driesche et al. 2019). Furthermore, a similar strategy has been successfully implemented for other RNA-binding proteins, including NOVA2 (Saito et al. 2019) and PABPC1 (Hwang et al. 2017; Jereb et al. 2018).

In order to study the dynamics of FMRP-RNA regulation rapidly after CA1-specific depolarization, we developed Opto-CLIP to allow for cell-type specific optogenetic activation of excitatory neurons in the mouse hippocampus (Fig. 1). Opto-CLIP incorporates rapid UV-irradiation after activation, allowing identification of RNAs directly crosslinked to FMRP precisely at the time of irradiation. To optogenetically manipulate neurons prior to CLIP or RiboTag, we performed bilateral stereotaxic injections of adeno-associated viruses (AAVs) to target the hippocampus (Fig. 2A). *Camk2a*-Cre mice were injected with either a Cre-inducible AAV which expresses the ChR2-mCherry fusion protein (Opto) or a Cre-inducible AAV which expresses a tdTomato reporter with no opsin (Control). Qualitative immunofluorescence (IF) confirmed that ChR2-mCherry expression was Cre-dependent (Fig. 2B) and restricted to CA1 neurons (Fig. 2C,D). We empirically determined that waiting 3-weeks after AAV-injection achieved optimal expression of the ChR2 opsin (Fig. 2C,D). Based on this timeframe, we sought to identify the ideal age of injection to restrict AAVs and molecular tags to area CA1 neurons. Consistent with previous reports (Sawicka et al. 2019; Hale et al. 2021; Tsien et al. 1996), we found that injecting mice at 8-weeks of age and sacrificing them for downstream molecular experiments at 11-weeks of age restricted *Camk2a*-Cre expression to CA1 neurons (Fig. 2C,D). AAV injections at later ages (> 8-weeks) led to a more-widespread expression of Cre-reporters in CA3 neurons and the dentate gyrus (DG) (Fig. 2E).

**Figure 1.**
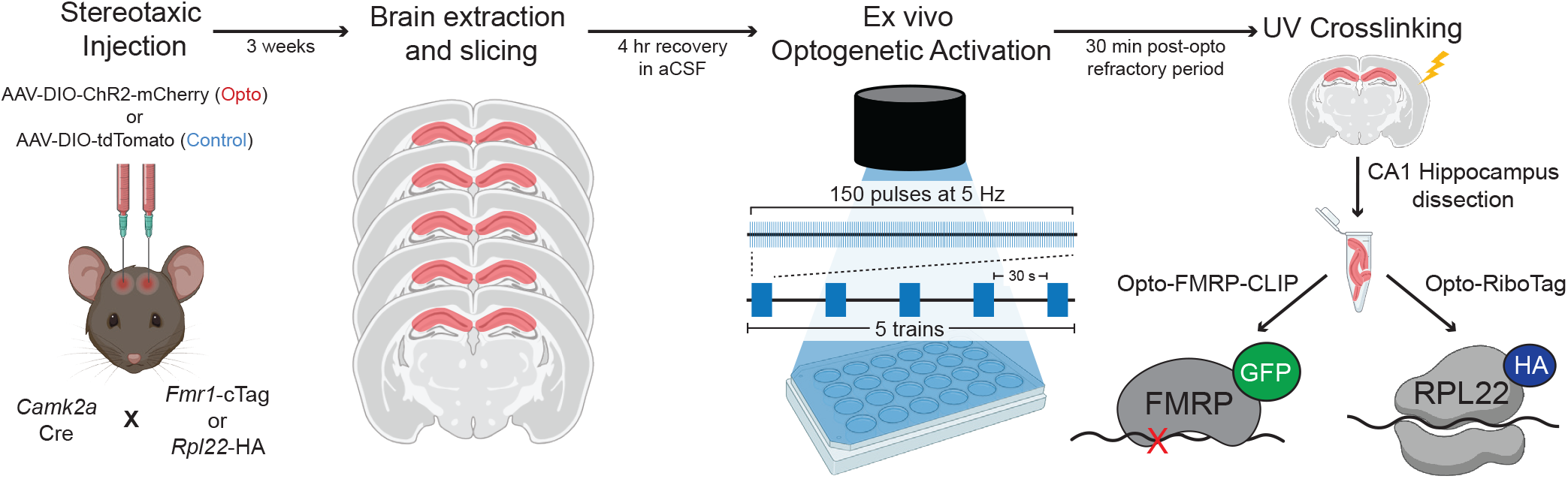
Overview of Opto-CLIP methodology. *Camk2a*-Cre mice are crossed to either *Fmr1*-cTag mice or *Rpl22*-HA (RiboTag) mice. 8-week-old *Camk2a*-Cre;*Fmr1*-cTag (for FMRP-CLIP) or *Camk2a*-Cre;*Rpl22*-HA (for RiboTag) are stereotaxically injected with either AAV-DIO-ChR2-mCherry (Opto) or AAV-DIO-tdTomato (Control) to enable Cre-dependent expression of ChR2 (H134R) fused to mCherry or tdTomato with no opsin, respectively, in CA1 excitatory hippocampal neurons. 3-weeks post-injection, mice are sacrificed and acute brain slices are pre-pared. After 4 hour recovery in aCSF, all slices are exposed to an LED stimulus protocol (blue light, 5 trains of 150 pulses at 5 Hz; 30” each). Slices are then recovered for 30 minutes and UV crosslinked (CLIP). CA1 hippocampus is then dissected and samples are assayed by FMRP-CLIP or RiboTag. Figure created with BioRender.com.

**Figure 2.**
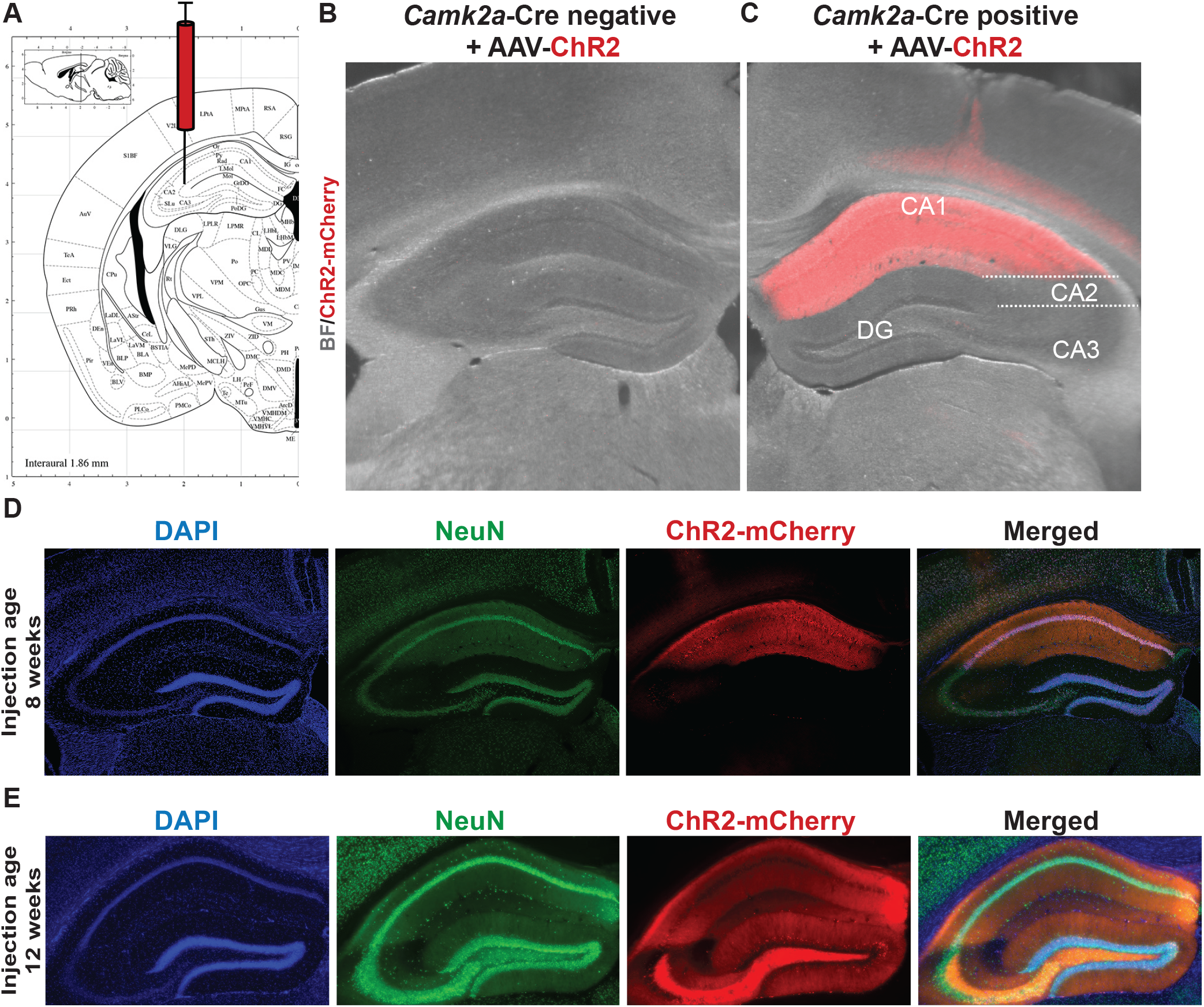
Cre-dependent expression of AAV-ChR2 in excitatory CA1 neurons of the adult mouse hippocampus. A) Schematic from Allen Brain Atlas showing region of the hippocampus targeted with AAVs. Opto mice were injected with pAAV-EF1a-DIO-hChR2-mCherry-WPRE-HGHpA. Control mice were injected with pAAV9-CAG-FLEX-tdTomato. Stereotaxic coordinates used (mm): Anterior/Posterior: -2, Medial/Lateral: +/-2, Dorsal/Ventral: -2. B and C) Overlaid Brightfield (gray) and fluorescent mCherry (red) images of the mouse hippocampus taken from *Camk2a*-Cre negative (B) and *Camk2a*-Cre positive (C) mice 3-weeks after stereotaxic injection of AAV-ChR2-mCherry. D and E) IF to detect AAV-mCherry expression in the mouse hippocampus 3-weeks after stereotaxic injection of AAV-mCherry. DAPI (blue) marker for nuclei, AAV-ChR2-mCherry (red), and NeuN (green) to label all neurons. Injections took place at 8-weeks of age (D) or 12-weeks of age (E). Magnification, 4X.

In order to assess neuronal health following AAV injection and acute slice preparation, we examined the intrinsic electrophysiological properties of CA1 pyramidal neurons. We injected *Camk2a*-Cre mice with AAV-ChR2-mCherry and performed fluorescence-guided whole-cell current clamp recordings in ChR2 positive neurons (Fig. 3A,B). We observed voltage decreases in response to a -50 pA and -25 pA hyperpolarizing current injections (Fig. 3C,D). Rheobase current, the minimal current to elicit an action potential (AP), was determined to be 50 pA in ChR2 positive CA1 neurons (Fig. 3C,D,E). Depolarizing current injections greater than 50 pA resulted in repetitive burst firing (Fig. 3C,D,E). AP frequency-current curves generated by the stepwise current injections revealed that ChR2 positive neurons increase AP firing in response to current injections and plateau at sustained AP frequencies upon 175 pA current injection (Fig. 3E). In order to further assess the integrity of neurons and their response to various ChR2 pulse frequencies, we performed whole cell recordings on ChR2 positive and ChR2 negative neurons during exposure to blue light. ChR2 positive, but not ChR2 negative, neurons displayed time-locked AP firing in response to blue light pulses at 1 Hz (Fig. 3F), 5 Hz (Fig. 3G), and 10 Hz (Fig. 3H) frequencies. All three frequencies successfully induced a single AP per light pulse for the entire duration of the stimulation protocols (10 seconds); however, 5 Hz was selected for the stimulation paradigm to avoid reduced membrane depolarization with later light pulses (Lin 2011), as seen with 10 Hz stimulation (Fig. 3H). Taken together, these data demonstrate that Cre-dependent ChR2 expression in CA1 neurons allows for optogenetic manipulation and establishes a paradigm to study cell-type specific RNA regulation following neuronal activation.

**Figure 3.**
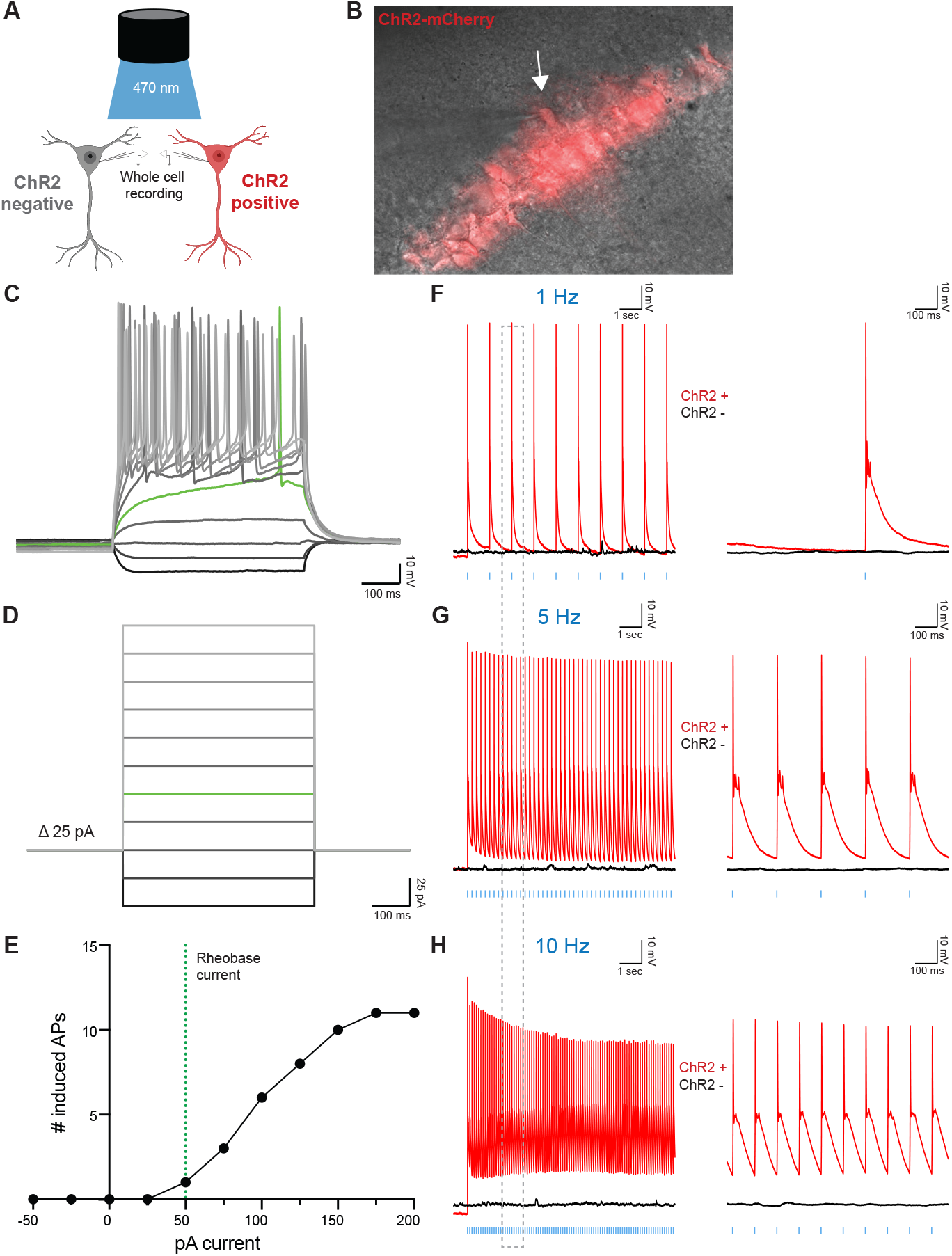
Optogenetic stimulation evokes action potentials in ChR2-expressing hippocampal excitatory neurons. A) Schematic of whole-cell configuration used for recordings of ChR2+ (visualized with red fluorescence) and ChR2-neurons. Figure created with BioRender.com. B) Overlaid Brightfield (gray) and fluorescent mCherry (red) microscope image showing ChR2+ CA1 neuron targeted for whole-cell patch clamping. Glass pipette tip and white arrow points to ChR2+ cell that underwent patch clamping. Magnification, 40X. C and D) Electrophysiological properties of ChR2+ neurons determined via stepwise injection of -50-200 pA current in 25 pA increments. Green lines show Rheobase current (50 pA), the minimal current to elicit an action potential (AP). Scale bars: 10 mV by 1 s. E) The number of AP induced by increasing injected currents. F-H) Whole-cell patchclamp traces of ChR2+ (red lines) or ChR2-(black lines) neurons stimulated with 10 sec of blue LED light at 1 Hz (F), 5 Hz (G), or 10 Hz (H). Vertical blue dashes indicate LED pulses. Right panels are 1 sec zoomed-in regions of left panels corresponding to the dotted gray box. Scale bars: 10 mV by 100 ms.

### Opto-CLIP identifies FMRP targets in control and activated neurons

A key innovation of using optogenetics as an activation paradigm to study FMRP-mediated RNA regulation is that both optogenetic activation and molecular analyses are done in a cell-type specific manner. Specifically, *Camk2a*-Cre activity drives expression of ChR2 and either HA-tagged ribosomes or GFP-tagged FMRP; the same neurons that are optogenetically activated can be molecularly assayed without the need for physical isolation, which could alter the RBP-RNA interactions.

To define FMRP-binding maps in optogenetically activated CA1 neurons, we modified the standard electrophysiology protocol (Fig. 3) to enable parallel processing of large sample sizes required for molecular assays. *Camk2a*-Cre;*Fmr1*-cTag mice were injected with either AAV-Control (Fig. 4A) or AAV-ChR2 (Fig. 4B) and 3 weeks later acute brain slices were prepared and allowed to recover for four hours to promote the return to baseline gene expression levels. We then modified the 10 second, 5 Hz optogenetic stimulation protocol mentioned above (Fig. 3G) to more closely mimic a protocol previously shown to induce hippocampal LTP (Kuwajima et al. 2020). Slices were therefore exposed to 5 trains of 30 second, 5 Hz pulses, recovered for 30 minutes, and then UV-crosslinked. We selected 30 minutes post-optogenetic stimulation to maintain consistency with prior studies of RNA changes after neuronal activation (Hacisuleyman et al. 2024).

**Figure 4.**
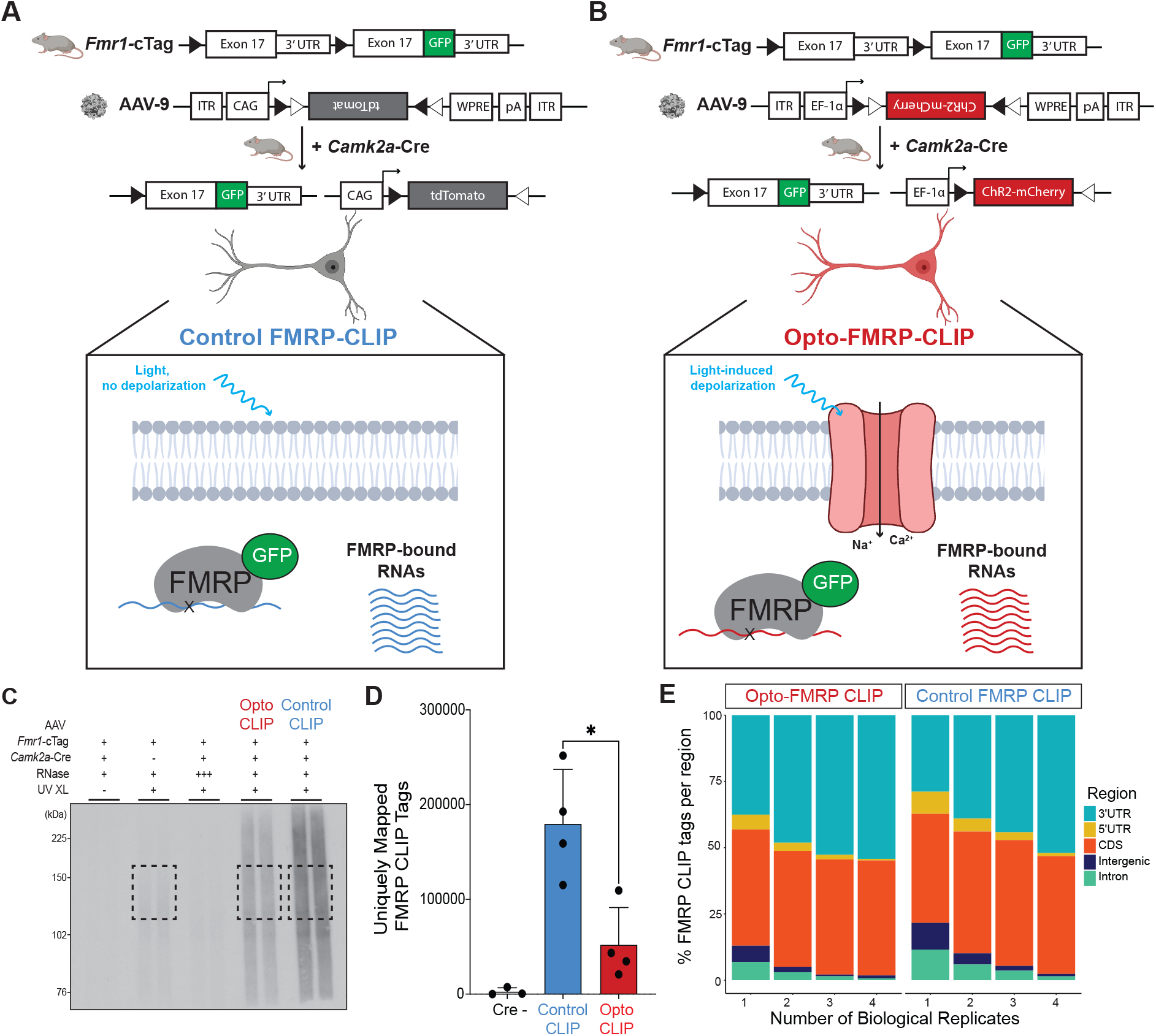
Opto-FMRP-CLIP uncovers RNAs differentially bound by FMRP in optogenetically activated neurons. A and B) Schematic depicting genetic targeting strategy for Control FMRP-CLIP (A) and Opto-FMRP-CLIP (B). Crossing *Fmr1*-cTag mice with *Camk2a*-Cre mice enables cell-type specific expression of GFP-tagged FMRP in excitatory neurons of the hippocampus. 3-weeks after stereotaxic injection of AAV-Control (A) or AAV-ChR2 (B), acute brain slices are exposed to LED activation paradigms which induces cell-type specific depolarization followed by UV crosslinking and molecular analysis by FMRP-CLIP. Figure created with BioRender.com. C) Autoradiograph image from Control and Opto-CLIP. Negative control samples consist of no UV Crosslinking (XL) (lane 1), Cre negative (lane 2), and high RNase treatment (lane 3). Black boxes indicate regions (120-175 kDa) subjected to CLIP sequencing. D) Comparison of uniquely mapped FMRP-CLIP tags obtained in replicate Control and Opto-CLIP experiments. See Methods for details of computational methods used to obtain uniquely mapped tags. Significance was calculated using a two-tailed, unpaired Student’s t-test. * p-value < 0.05. n=3, Cre negative samples. n=4 for Control and Opto-CLIP samples. E) Genomic annotations of CLIP tags separated by biological complexity (BC).

To isolate FMRP-bound RNA, crosslinked samples were immunoprecipitated with anti-GFP antibodies, and RNA fragments were isolated, stringently purified, and sequenced using previously defined CLIP protocols (Hwang et al. 2017; Sawicka et al. 2019; Hale et al. 2021) (Fig. 4C). Cre negative samples were used as a negative control and processed in parallel (Fig. 4C, second lane). Reads were mapped to the transcriptome and the number of uniquely mapped tags (UMTs) was determined for each replicate. Cre negative samples had very few UMTs compared to Cre positive samples (Fig. 4D), demonstrating the low signal to noise of FMRP-CLIP. Interestingly, we observed a significant decrease (∼3.5-fold) in FMRP binding in light-treated neurons compared to controls (Fig. 4D). Principal component analysis (PCA) revealed that Opto-CLIP and Control samples clustered as distinct groups (Supplemental Fig. 1A) and samples were highly correlated across replicates (Supplemental Fig. S1B). Mapping the genomic distribution of tags with increasing Biological Complexity (BC) (requiring a tag to be found in “n” biological replicates) revealed a preference for CDS and 3’ UTRs, with Opto-CLIP and Control samples having similar distributions (Fig. 4E), consistent with prior observations of FMRP-CLIP in mouse brain (Darnell et al. 2011; Sawicka et al. 2019; Hale et al. 2021). In total, 3661 genes contained at least 5 FMRP CLIP tags in all four Opto-CLIP or Control experiments (BC = 4). Taken together, Opto-FMRP-CLIP accurately defined RNAs bound by FMRP in activated neurons.

### Opto-RiboTag identifies ribosome-associated transcripts in control and activated neurons

For several well-studied RBPs, quantification of differential binding between two conditions has been accomplished by calling peaks from CLIP tags and then qualitatively comparing tags in peaks across conditions (Hwang et al. 2017; Weyn-Vanhentenryck et al. 2014; Moore et al. 2018). FMRP, however, has been shown to display continuous coverage across the coding sequence of its targets (Darnell et al. 2011; Sawicka et al. 2019; Hale et al. 2021), and consequently traditional peak-calling algorithms fail to capture the nature of FMRP binding to its target transcripts. As an alternative, studies have mapped FMRP CLIP tags to the transcriptome and normalized FMRP binding relative to transcript abundance in that same cell type (Sawicka et al. 2019; Hale et al. 2021). To measure transcript abundance, we crossed *Camk2a*-Cre mice with the RiboTag mouse (Sanz et al. 2009), enabling Cre-mediated HA tagging of all ribosome bound transcripts in CA1 neurons. Immunoprecipitation (IP) with an HA antibody, in the presence of cycloheximide to maintain ribosome-mRNA association, enables isolation of all ribosome-bound transcripts. Although this strategy limits quantitation to ribosome-bound RNAs, FMRP is predominantly associated with polyribosomes in the brain (Darnell et al. 2011), and in-depth analyses have demonstrated cell-type specific RiboTag efficiently isolates the majority of the transcriptome (Sawicka et al. 2019; Hale et al. 2021; Heiman et al. 2008, 2014; Doyle et al. 2008).

To define ribosome associated transcripts in activated CA1 neurons, we injected *Camk2a*-Cre;*Rpl22*-HA mice with either AAV-Control (Fig. 5A) or AAV-ChR2 (Fig. 5B) and performed Control and Opto-RiboTag. Brain slices were optically activated and area CA1 was dissected to enrich for CA1 neurons and deplete HA-expressing neurons from CA3 and DG (Fig. 5C, white dotted region), as for Opto-CLIP, excluding the crosslinking step. Rpl22-HA positive, *Camk2a*-Cre negative mice were used as controls and processed in parallel (Fig. 5D; right three lanes; Fig. 5E; left column). We optimized the concentration of HA antibody used to maximize HA-depletion post-IP (Fig. 5D). We detected similar levels of HA protein in ChR2 and Control samples and HA protein was absent in Cre negative samples (Fig. 5D). Cre positive samples, but not Cre negative samples, had enrichment of CA1 specific excitatory neuronal genes (*Neurod6, Camk2a, Rbfox3, Snap25, Nrgn, Hpca, Crym*, and *Chn1*), while markers of other cell types, such as inhibitory neurons (*Gad2, Sst*, and *Calb2*), oligodendrocyte precursor cells (OPCs) (*Pdgfra* and *Ptprz1*), oligodendrocytes (*Mag, Mal, Mpb, Mobp*, and *Plp1*), and astrocytes (*Gfap, Glul, Aqp4, Aldh1l1, Pla2g7, Slc1a3*, and *Aldoc*) were depleted (Fig. 5E). These data demonstrate that mRNAs isolated by RiboTag IP originate from CA1 excitatory neurons.

**Figure 5.**
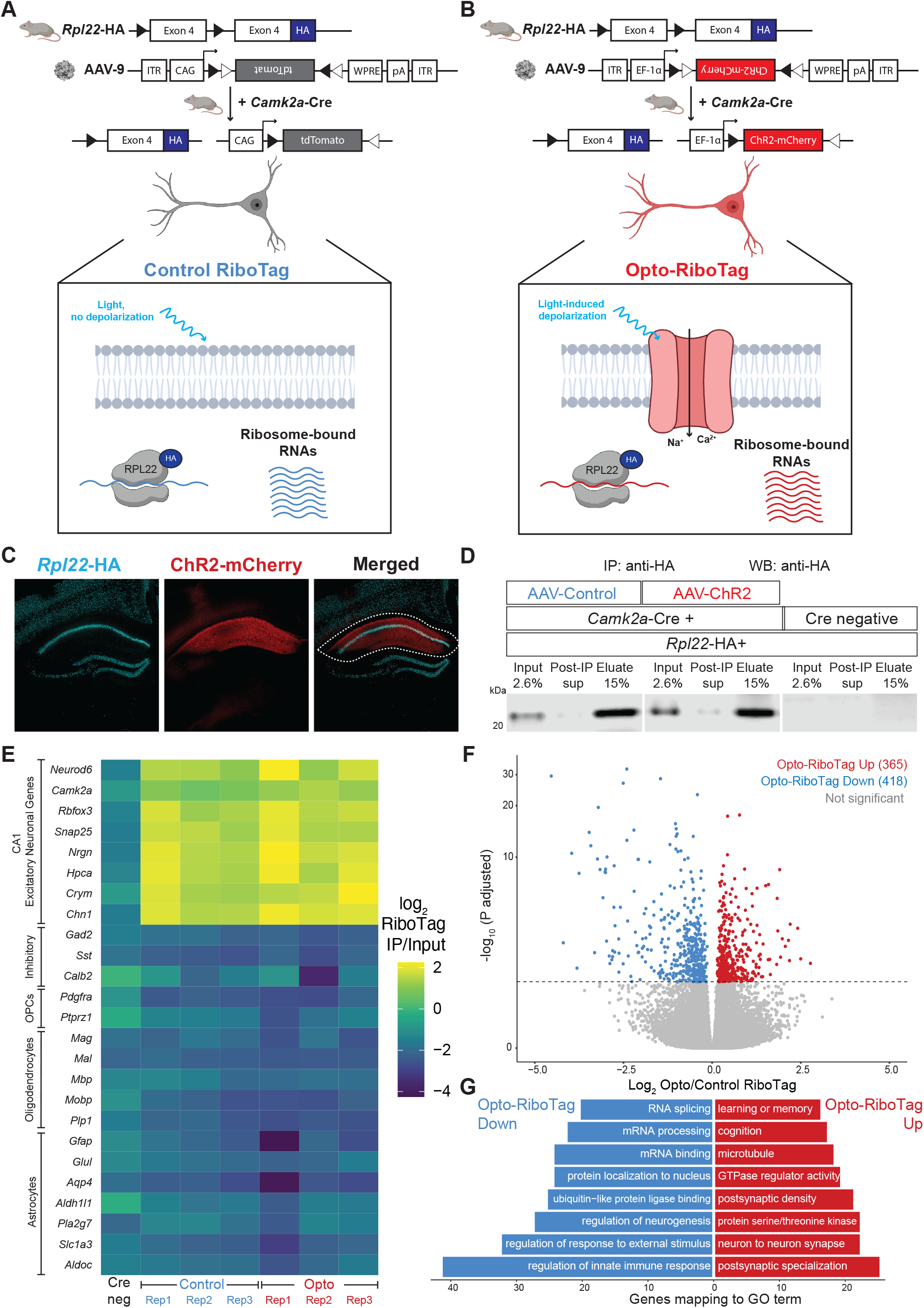
Opto-RiboTag robustly defines ribosome-bound RNA in optogenetically activated and control neurons. A and B) Schematic depicting genetic targeting strategy for Control RiboTag (A) and Opto RiboTag (B). Crossing *Rpl22*-HA mice with *Camk2a*-Cre mice enables cell-type specific expression of HA-tagged ribosomes in excitatory neurons of the hippocampus. 3-weeks after stereotaxic injection of AAV-Control (A) or AAV-ChR2 (B), acute brain slices are exposed to LED activation paradigms which induces cell-type specific depolarization and molecular analysis is performed by HA-IP and RNA-sequencing of input and immunoprecipitated RNA. Figure created with BioRender.com. C) Microscope images showing IF done on brain slices from *Camk2a*-Cre;*Rpl22*-HA mice 3-weeks following stereotaxic injection of AAV-mCherry. DAPI (blue) marker for nuclei, AAV-mCherry (red), and HA (cyan) to label *Camk2a*-Cre responsive neurons that will undergo optogenetic activation and molecular analysis by RiboTag. White dotted line indicates the CA1 hippocampus that is dissected prior to RiboTag. Magnification, 10X. D) Western blot for HA shows that HA protein is restricted to Camk2a-Cre+ samples (left 6 lanes), is sufficiently depleted from cell lysate by immunoprecipitation (IP) with HA-antibody (Post-IP sup lanes), and is enriched in eluate following HA-IP. Marker showing size of 20 kDa ladder. E) Heatmap of the RiboTag enrichment scores calculated by log_2_ (RiboTag IP Transcript Per Million (TPM)/RiboTag Input TPM) following HA-immunoprecipitation from Cre negative and Cre positive RiboTag samples. Yellow color indicates positive enrichment of CA1 excitatory neuronal genes and dark blue color indicates negative enrichment of non-excitatory markers, such as inhibitory, OPC (oligodendrocyte precursor cells), oligodendrocytes, and astrocytes genes. Cre negative sample is from Cre negative;*Rpl22*-HA mice subjected to the same RiboTag pipeline as the other samples. All CA1 genes listed are significantly enriched in Cre-positive IP samples versus Input samples (p.adjusted < 0.05; log_2_ fold change IP/Input > 0). All other genes listed are significantly depleted in Cre-positive IP samples versus Input samples (p.adjusted < 0.05; log_2_ fold change IP/Input < 0) Significance was calculated from DESeq2 with correction for multiple testing correction done via the Benjamini–Hochberg method. F) Volcano plots showing RNA transcripts upregulated (red) and downregulated (blue) in Opto RiboTag versus Control RiboTag. Transcripts are considered significant with a p.adjusted < 0.05. Significance was calculated from DESeq2 with correction for multiple testing correction done via the Benjamini–Hochberg method. G) Gene Ontology (GO) analysis on ribosome bound RNAs downregulated (blue/ left) or upregulated (red/right) following optogenetic activation. Counts indicate the number of genes found in each category. All categories listed are p.adjusted (Benjamini–Hochberg) < 0.05 measured by enrichGO (Yu et al. 2012).

Sequencing analyses of Control and Opto-RiboTag samples uncovered RNAs differentially bound by ribosomes upon activation, of which 365 transcripts were upregulated and 418 transcripts were downregulated 30 minutes following optogenetic activation (Fig. 5F). To explore the pathways involved in these molecular changes, we performed gene set enrichment analysis (GSEA) on upregulated and downregulated transcripts from Control and Opto-RiboTag. Interestingly, gene sets associated with learning and memory, cognition, and postsynaptic density were enriched in transcripts upregulated in Opto-RiboTag (Fig. 5G, red bars). In contrast, downregulated transcripts encode proteins involved in RNA splicing, mRNA processing, and protein localization to nucleus (Fig. 5G, blue bars). This indicates that different biologically coherent sets of ribosome-bound transcripts–those involved in synaptic plasticity and higher order levels of RNA regulation, respectively– are preferentially translated after depolarization. Taken together, these data demonstrate that Opto-RiboTag is a sensitive technique for uncovering cell-type specific changes in RNA regulation following neuronal activation.

### FMRP targets are dynamically regulated in activated neurons

We next sought to integrate our findings from Opto-RiboTag and Opto-FMRP-CLIP to build a model of FMRP-mediated RNA regulation in activated neurons. Previous studies demonstrated that FMRP CLIP tag density on a specific transcript is significantly correlated with the abundance of the transcript (Sawicka et al. 2019; Hale et al. 2021; Doyle et al. 2008); however, analyses were limited to steady state conditions. To determine if this finding holds true in activated neurons, we compared transcript abundance in Opto-RiboTag and Opto-FMRP-CLIP samples. Parallel analyses were done to compare Control RiboTag to Control FMRP-CLIP. We found that transcript abundance measured by RiboTag and CLIP were significantly positively correlated in both control and activated conditions (Supplemental Fig. S2A-H). We used this relationship to calculate an FMRP CLIP score for each transcript that quantifies the amount of FMRP binding to a transcript relative to other transcripts of similar abundance. We compared average Control or Opto-RiboTag (log_2_ TPM) to each individual Control or Opto-CLIP (log_2_ TPM) replicate and fit each plot with a linear regression line (Supplemental Fig. S2A-H). For each transcript, we calculated Control and Opto-CLIP scores so that the higher the score, the greater the FMRP binding relative to similarly abundant transcripts. For each condition, FMRP binding stringency was calculated by averaging Control and Opto-CLIP scores across replicates. In control neurons, there were 1602 stringent FMRP targets (Fig. 6A) compared to 1027 stringent targets in activated neurons (Fig. 6B). To determine how FMRP binding changed with optogenetic activation, we further classified all transcripts based on Control and/or Opto-FMRP-CLIP scores. 898 transcripts had stringent CLIP scores in both activated and control conditions (Fig. 6C, green dots), 129 transcripts had stringent CLIP scores only in activated conditions (Fig. 6C, red dots), 704 transcripts had stringent CLIP scores only in control conditions (Fig. 6C, blue dots), and most (10436) transcripts showed relatively low levels of FMRP binding in both conditions (Fig. 6C, gray dots). These data identify stringently defined FMRP bound and unbound transcripts in activated and control neurons.

**Figure 6.**
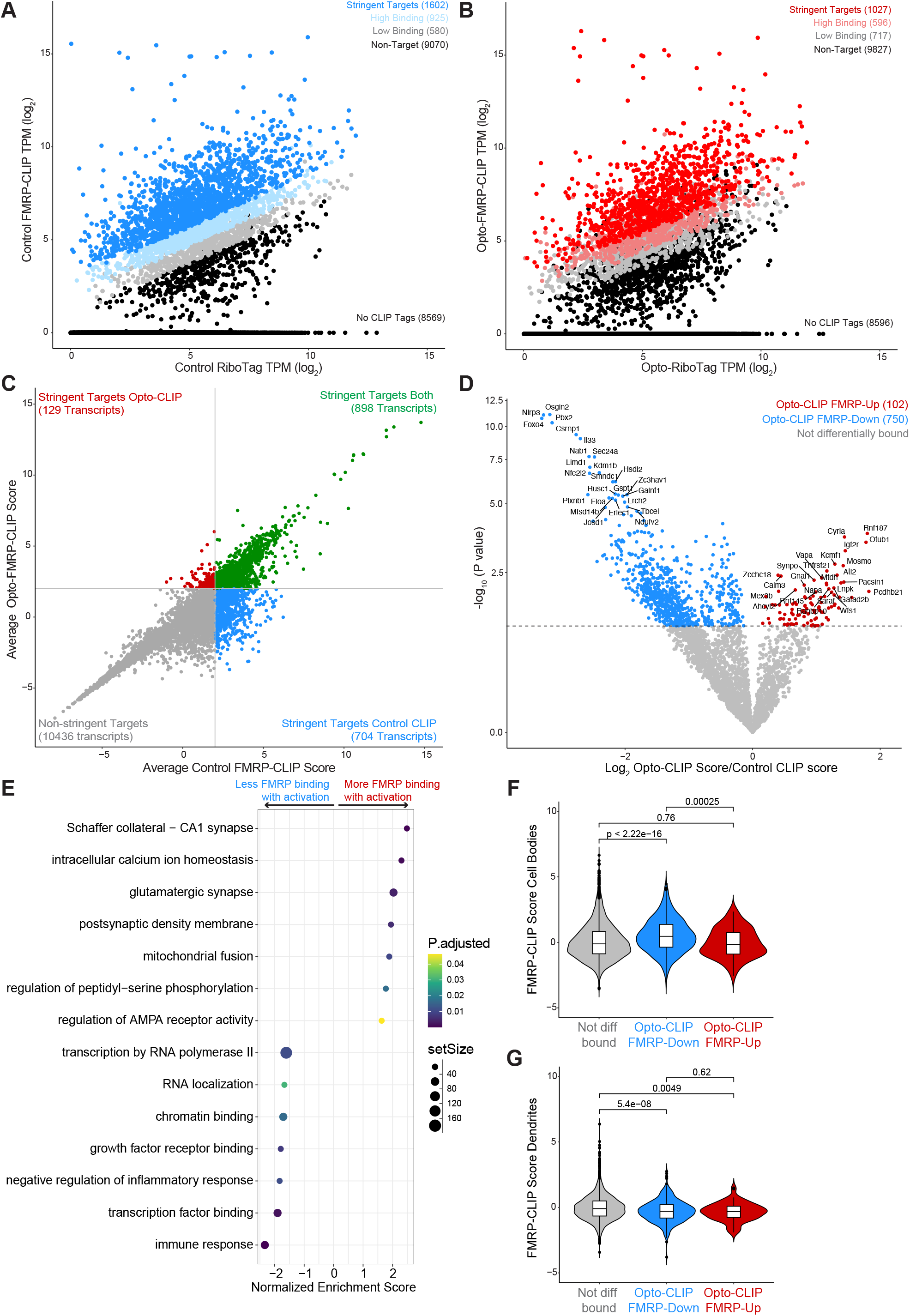
Neuronal activation alters FMRP binding to transcripts essential for neuronal function. A and B) Scatter plots of log_2_ normalized transcript per million (TPM) of CLIP tags from averaged Control (A) or Opto-CLIP (B) experiments compared to average log_2_ RiboTag TPM from Control (A) or Opto-RiboTag (B). n=4 biological replicates for FMRP-CLIP. n=3 biological replicates for RiboTag. Stringent targets were defined as those with an average CLIP score > 2, targets with high binding were defined as those with an average CLIP score > 1 and targets with low binding were defined as those with an average CLIP score between 0 and 1. Targets of each subclass are highlighted in the plot and the number of genes within each subclass is indicated. See methods for further details on CLIP score calculations. C) Scatter plot of Opto-CLIP scores compared to Control CLIP scores for each transcript. Green dots indicate transcripts with CLIP scores > 2 in both conditions. Blue dots indicate transcripts with CLIP scores > 2 in control neurons and < 2 in activated neurons. Red dots indicate transcripts with CLIP scores < 2 in control neurons and > 2 in activated neurons. Gray dots indicate transcripts with CLIP scores < 2 in both conditions. D) Volcano plot comparing log2 fold changes and p-values comparing Control versus Opto-CLIP scores (significance determined by Limma (Ritchie et al. 2015), p-value < 0.05). Transcripts with significantly higher Opto-CLIP scores are shown in red (FMRP-Up). Transcripts with significantly lower Opto-CLIP scores are shown in blue (FMRP-Down). For each category, the top 25 most statistically significant transcripts are labeled. E) Gene Set Enrichment Analysis (GSEA) on transcripts differentially bound by FMRP as shown in D. Positive normalized enrichment score (NES) indicates transcripts more bound by FMRP in activated conditions (red dots in D) and negative NES indicates transcripts less bound by FMRP in activated conditions (blue dots in D). F and G) Violin boxplots showing FMRP CLIP scores calculated from microdissected CA1 cell bodies (F) and dendrites (G) (Hale et al. 2021). Red boxes show transcripts with significantly higher Opto-CLIP scores. Blue boxes show transcripts with significantly lower Opto-CLIP scores. Wilcoxon rank-sum test was used to determine p-values indicated.

To quantify the change in FMRP binding after activation, we compared CLIP scores in control versus activated neurons for each transcript (Fig 6D). We discovered that the vast majority (84%; 750/852) of differentially bound targets were less bound by FMRP in activated versus control neurons (Fig. 6D, blue dots, herein referred to as “FMRP-Down”). Only 14% (102/852) of transcripts exhibited more FMRP binding in activated neurons compared to control neurons (Fig. 6D, red dots, herein referred to as “FMRP-Up”). Interestingly, when examined by GSEA, “FMRP-Up” transcripts were enriched in synaptic functions, such as regulation of AMPA receptor activity, Schaffer collateral—CA1 synapse, glutamatergic synapses, and intracellular calcium ion homeostasis (Fig. 6E). In contrast, “FMRP-down” transcripts were associated with nuclear functions, such as chromatin binding, transcription factor binding, transcription by RNA polymerase II, and RNA localization (Fig. 6E).

These findings were reminiscent of prior data indicating that FMRP has distinct RNA binding preferences in subcellular regions of CA1 neurons (Hale et al. 2021). Hence, we compared FMRP CLIP scores in transcripts previously identified as FMRP cell body or synaptic targets. Remarkably, we found that we could distinguish these two transcript sets—those encoding synaptic targets (FMRP-Up) had significantly lower cell body CLIP scores than “FMRP-Down” transcripts associated with nuclear functions (Fig. 6F). We further observed that cell-body associated FMRP regulated (FMRP-Down) transcripts had significantly lower dendritic CLIP scores than transcripts with no activity-induced changes in FMRP binding (Fig. 6G). Importantly, these observations were in agreement with GSEA terms associated with FMRP-Up and FMRP-Down transcripts (Fig. 6E). These findings demonstrate that after activation of CA1 neurons, FMRP is less bound to transcripts in the cell body, which are involved in nuclear RNA regulatory processes, but shows little change in transcripts encoding synaptic targets, which are present at the synapse. Taken together, these data suggest the possibility that there is differential regulation of FMRP binding in the cell body subcellular compartment after depolarization, and is discussed below.

We observed that most (84%) differentially bound FMRP targets—primarily those present in the cell body encoding nuclear processes— showed reduced binding in activated neurons (Fig. 6D,E,F). This observation led us to hypothesize that FMRP may be preferentially released from these transcripts upon neuronal activation, potentially freeing them from ribosome stalling. Given previous work showing FMRP acts as a reversible translational repressor (Darnell et al. 2011; Sawicka et al. 2019; Hale et al. 2021), we also explored whether, in the 30 minutes between optogenetic activation and CLIP, some of these transcripts displayed differential ribosome binding—ranging from ribosome runoff to secondary ribosome re-engagement.

To investigate this, we overlaid differential FMRP-CLIP binding data with transcript abundance measured by RiboTag for the 750 transcripts that were less bound by FMRP in activated neurons (FMRP-Down). Of these, 72 transcripts showed decreased ribosome binding after activation (Fig. 7A, orange dots), while 94 transcripts had increased ribosome association (Fig. 7A, violet dots). A minority of transcripts (102) had increased FMRP binding in activated neurons (FMRP-Up); of these, 9 had increased ribosome association (Supplemental Fig. 3A, green dots), 10 transcripts had decreased ribosome binding (Supplemental Fig. 3A, magenta dots), and the majority showed no change. In sum, these data suggest that within the cell body, FMRP-regulated transcripts may be preferentially “translationally de-repressed”, as evidenced by less FMRP binding and altered ribosome binding following neuronal activation.

**Figure 7.**
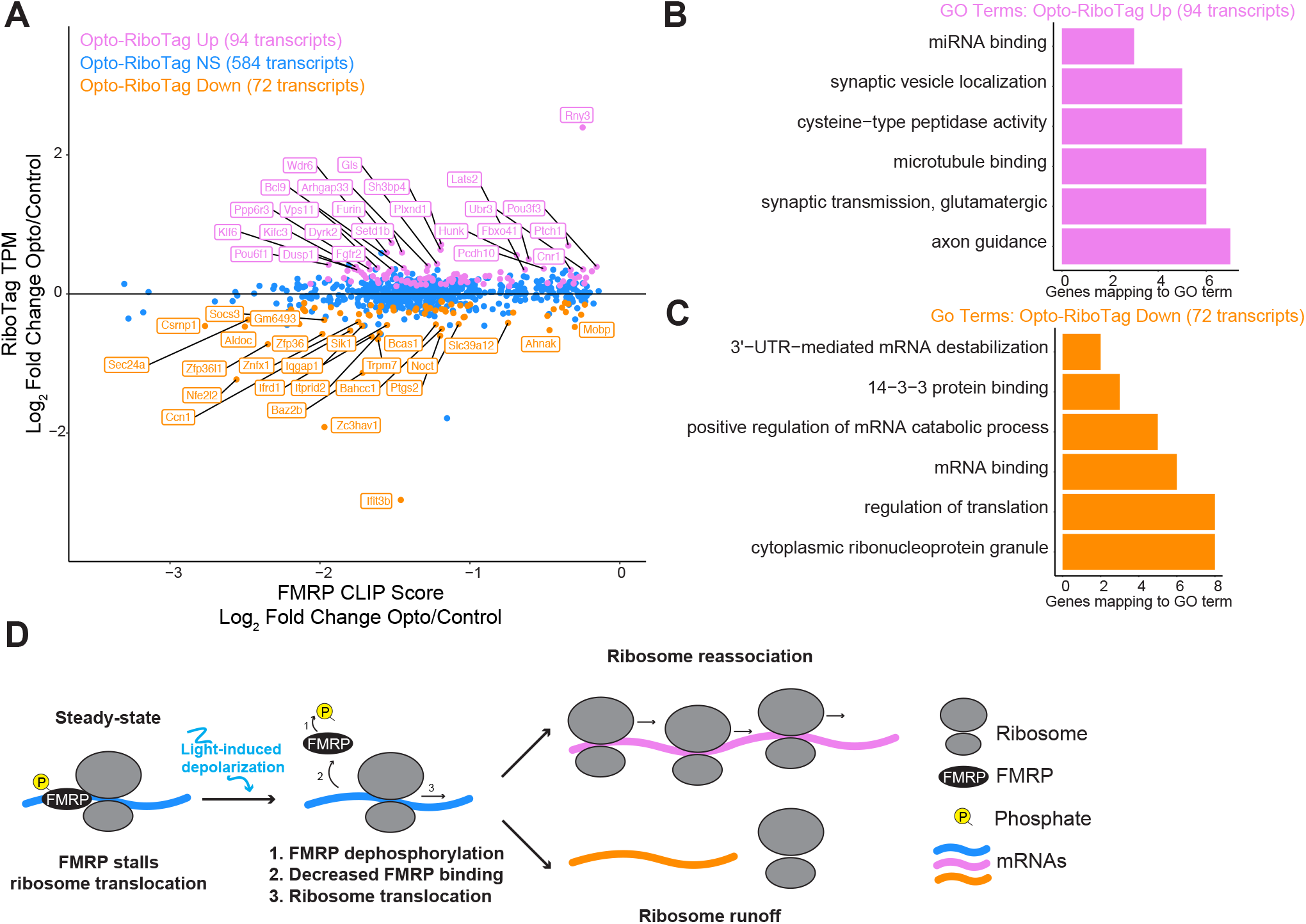
Reduced FMRP binding in activated neurons triggers dynamic regulation of functionally distinct transcripts. A) Scatter plot of Opto-RiboTag versus Control RiboTag (log_2_ TPM) compared to Opto-CLIP versus Control CLIP CLIP scores (log_2_ fold change). Blue dots indicate transcripts that are less bound by FMRP in activated neurons (FMRP-Down) and are changed by Opto-RiboTag. Violet dots indicate transcripts that are less bound by FMRP in activated neurons (FMRP-Down) and are upregulated in activated RiboTag. Orange dots indicate transcripts that are less bound by FMRP in activated neurons (FMRP-Down) and are downregulated in activated RiboTag. 50 transcripts with largest RiboTag log_2_ fold changes are labeled. B) GO analysis on transcripts that are less bound by FMRP in activated neurons (FMRP-Down) and are upregulated in activated RiboTag corresponding to the Violet dots in A. C). GO analysis on transcripts that are less bound by FMRP in activated neurons (FMRP-Down) and are downregulated in activated RiboTag corresponding to the Orange dots in A. Bar length corresponds to the number of genes mapping to a GO term. All categories listed are p.adjusted (Benjamini–Hochberg) < 0.05 measured by enrichGO (Yu et al. 2012). D) Model depicting FMRP regulation of distinct subsets of transcripts in optogenetically activated neurons. In steady-state neurons, FMRP binds to mRNAs and represses translation by stalling ribosome translocation (Darnell et al. 2011). Upon neuronal activation, such as optogenetic stimulation in this study, FMRP undergoes calcium-dependent dephosphorylation (Bear et al. 2004; Lee et al. 2011) and decreases binding to target transcripts. For a subset of transcripts (violet mRNAs), reduced FMRP binding triggers increased transcript abundance measured by RiboTag, likely due increased ribosome reassociation. In contrast, for another subset of transcripts (orange mRNAs), reduced FMRP binding leads to decreased transcript abundance measured by RiboTag, potentially due to ribosome runoff.

To explore functional differences between transcripts with reduced FMRP binding after activation that also displayed differential ribosome association, we performed GSEA on these transcripts (Fig. 7A, violet and orange dots). Transcripts that were more ribosome-associated and less bound by FMRP in activated neurons were significantly enriched in pathways related to miRNA binding, synaptic vesicle localization, microtubule binding, and synaptic transmission (Fig. 7B). In contrast, transcripts that were less ribosome-associated and less bound by FMRP were enriched in pathways involving ribonucleoprotein granules, translational regulation, 14-3-3 protein binding, and 3’−UTR−mediated mRNA destabilization (Fig. 7C). Taken together, these findings suggest that neuronal activation induces changes in FMRP binding, leading to the up- and downregulation of functionally distinct classes of mRNAs. Furthermore, the dynamics of ribosome run-off and reassociation appear to differ between these transcript groups (Fig. 7D).

## DISCUSSION

FMRP is critical for neuronal processes, including synaptic plasticity, which underlies learning and memory (Bear et al. 2004; Huber et al. 2002; Lauterborn et al. 2007; Lee et al. 2011; Huber et al. 2000). However, the global dynamics of FMRP:RNA interactions during neuronal activation remain undefined. Here, we present Opto-CLIP, a platform used here to investigate cell-type-specific RNA regulation by FMRP in optogenetically activated CA1 excitatory neurons. Using Opto-CLIP, we identified significant changes in FMRP-binding patterns and ribosome-associated RNA profiles following neuronal activation.

We find that FMRP regulates two distinct subsets of transcripts. The predominant subset (∼86%; 750/853) showed decreased FMRP binding after optogenetic activation (Opto-CLIP FMRP-Down; Fig. 6D; blue dots) and is enriched for genes encoding nuclear processes, such as transcription and chromatin binding, which are preferentially bound by FMRP in the CA1 cell soma (Hale et al. 2021)‥ This suggests that in the 30 minute window after CA1 neuronal depolarization, FMRP translational repression is selectively relieved for transcripts regulating nuclear functions. In contrast, a smaller subset of transcripts (14%; 102/853) exhibited increased FMRP binding after activation (Opto-CLIP FMRP-Up; Fig. 6D; red dots), and these encode proteins involved in synaptic functions. These findings suggest distinct temporal dynamics for FMRP regulation of transcripts in the cell soma versus the synapse.

Among transcripts with reduced FMRP binding upon activation, RiboTag analysis revealed roughly equal numbers of transcripts that were upregulated and downregulated. This dual regulation is consistent with prior studies demonstrating that FMRP can regulate transcripts by promoting and repressing translation (Aryal and Klann 2018), as well as Translating Ribosome Affinity Purification (TRAP) studies comparing *Fmr1*-KO and WT CA1 hippocampi (Thomson et al. 2017). A plausible model is that FMRP is released from transcripts in the cell body within 30 minutes after depolarization, releasing its block on ribosomal elongation (Darnell et al. 2011) and enabling translation. Some transcripts may subsequently rebind FMRP, arresting newly bound ribosomes (violet transcripts in Fig. 7) and initiating another cycle of translational control. Others remain ribosome free after runoff (orange transcripts in Fig. 7), suggesting additional features of translational control.

FMRP likely exerts its effects on different transcripts or compartments at different time points. Our study focused on the 30 minute mark based on prior work detailing RNA changes after neuronal activation (Hacisuleyman et al. 2024). It is plausible that FMRP has both rapid and slower translational controls, depending on the transcript and its cellular context. Future experiments investigating FMRP-mediated RNA regulation across different timescales will be important for resolving these mechanisms.

Together, these findings support a model in which FMRP exerts differential effects in distinct subcellular compartments of the same CA1 neuron. In the soma, FMRP release facilitates translational derepression of transcripts involved in nuclear regulatory processes, positioning FMRP as a regulatory sensor of neuronal activation. This sensor triggers biologically coherent responses specific to the soma. At 30 minutes after depolarization, we observe less evidence of FMRP dynamics in transcripts that are associated with the synapse, possibly due to faster turnover or distinct regulation in this compartment. Overall, these results encourage further exploration of a model in which FMRP dampens synaptic signaling while enhancing nuclear transcriptional and RNA regulatory controls 30 minutes after depolarization. Such a model positions FMRP as a critical integrator linking synaptic activity to nuclear responses, thereby acting as a mediator of homeostatic plasticity (Darnell 2020; Turrigiano and Nelson 2004).

While our study provides significant advancements in understanding FMRP-mediated RNA regulation, there are limitations to consider. The optogenetic activation paradigm, though advantageous for its rapid and reversible nature, may not fully recapitulate the complexities of in vivo neuronal activation and synaptic plasticity. Additionally, while we demonstrated significant changes in FMRP binding and ribosome association upon neuronal activation, their functional implications require further exploration. Future studies integrating electrophysiology, protein analysis, and other neuronal subtypes or brain regions will provide a more comprehensive understanding of FMRP’s role in the nervous system. Importantly, Opto-CLIP offers a versatile platform for studying RBPs, with broad applications in investigating RNA regulation in neuronal and non-neuronal contexts.

In conclusion, we developed Opto-CLIP as a broadly applicable platform for studying cell-type-specific RNA regulation by RBPs in response to neuronal activity. By combining Opto-FMRP-CLIP and Opto-RiboTag, we constructed a comprehensive model of FMRP-mediated RNA regulation. These insights shed light on the molecular mechanisms underlying synaptic plasticity and cognitive functions and lay the foundation for future research into the molecular basis of learning, memory, and neurodevelopmental disorders.

## MATERIALS AND METHODS

### Mice

All animal procedures were approved by The Rockefeller University Institutional Animal Care and Use Committee (IACUC) and mice were treated in accordance with the National Institutes of Health Guide for the Care and Use of Laboratory Animals. Mice were allowed ad libitum access to food and water at all times, weaned at 3 weeks of age, and maintained on a 12-hour light/dark cycle. All mouse lines used in these studies were on the C57BL/6 genetic background. B6.Cg-Tg (*Camk2a*-cre)T29-1Stl/J (*Camk2a*-Cre) (Tsien et al. 1996), B6N.129-Rpl22tm1.1Psam/J (RiboTag)(Sanz et al. 2009) were purchased from Jackson Laboratories. The *Fmr1*-cTag mouse has been described and characterized elsewhere (Van Driesche et al. 2019). Briefly this mouse line was generated by introducing loxP sites either side of the terminal exon of the *Fmr1* gene followed by a downstream AcGFP-tagged version of the terminal exon and surrounding intronic sequences. Thus, either FMRP or AcGFP-tagged FMRP can be expressed from the cTag allele in a mutually exclusive manner, dependent on Cre expression. Further details about mouse strains used in this study can be found in Supplemental Table S1.

### Stereotaxic Surgeries

Stereotaxic surgeries were performed on 8-week-old *Camk2a*-Cre mice crossed to either *Fmr1*-cTag mice or *Rpl22*-HA mice. Mice were anesthetized with inhaled isoflurane (Kent Scientific; SomnoFlo) at 3% for induction. Once anesthetized, animals were placed in the stereotaxic frame (Kopf; Model 1900) and exposed to 0.25% isoflurane for the remainder of surgery. Mice were shaved and paralube (artificial tears) applied to the eyes to avoid drying. The surgical field was cleaned with betadine solution followed by 70% ethanol. A sterile scalpel was used to perform a midline incision to expose the skull. The position of bregma was determined and used as a landmark for stereotaxic coordinates. The following coordinates (in mm) were used for targeting CA1 neurons (Anterior/ Posterior: -2, Medial/Lateral: +/-2, Dorsal/Ventral: -2). Two small burr holes (0.6 mm) were drilled bilaterally over the target site of injection. A 34G Beveled needle (World Precision Instruments; NF34BV-2) attached to a 10 μl Hamilton syringe was used to inject one microliter of virus per site at a rate of 0.095 μl/min. Following virus injections, the needle was left in place for 5 minutes before being slowly retracted. For conditional expression of channelrhodopsin in the mouse hippocampus, mice were injected with pAAV-EF1a-double floxed-hChR2(H134R)-mCherry-WPRE-HGHpA (a gift from Karl Deisseroth; Addgene: 20297). For conditional expression of the control reporter, mice were injected with pAAV-FLEX-tdTomato (a gift from Edward Boyden; Addgene: 28306). Both plasmids were AAV9 serotypes and were diluted 1:20 which corresponded to 1.25 E+09 genomic particles per injection site. Further details about AAVs used in this study can be found in Supplemental Table S1. The scalp incision was closed with dissolvable surgical sutures (4-0 Coated Vicryl Violet 1×27” FS-2; Ethicon; J397H) and mice were given a subcutaneous injection of Meloxicam SR (6 mg/kg) for pain relief that lasts 48 hours. In addition, 1.0 mL sterile saline was administered subcutaneously to prevent further dehydration. Mice were allowed to recover from surgery for 3 weeks prior to any downstream experiments.

### Brain Slice Preparation

At 11 weeks of age, mice were deeply anesthetized with 3% inhaled isoflurane (Kent Scientific; SomnoFlo) and perfused transcardially with 10 mL ice cold dissection buffer (2.5 mM KCl, 0.5 mM CaCl2 dihydrate, 7 mM MgCl2 hexahydrate, 25 mM NaHCO3, 1.25 mM NaH2PO4, 11.6 mM Sodium L-ascorbate, 3.1 mM sodium pyruvate, 110 mM choline chloride, 25 mM glucose). Brains were sectioned at 400 μm on a Vibratome (Leica VT1200 S) and placed in the Brain Slice Keeper (AutoMate Scientific BSK-4) in artificial cerebrospinal fluid (aCSF) (2.5 mM KCl, 118 mM NaCl, 1.3 mM MgCl2, 2.5 mM CaCl2, 26 mM NaHCO3, 1 mM NaH2PO4, 10 mM glucose) constantly perfused with 95% O2 / 5% CO2 gas (carbogen). Slices were allowed to recover at 32°C for 1 hour. For all electrophysiological recordings, slices were allowed to recover for an additional 30 minutes at room temperature in carbogenated aCSF. For all CLIP and RiboTag experiments, slices recovered for an additional 3 hours at room temperature in carbogenated aCSF.

### Electrophysiological Recordings

Glass capillary pipettes were pulled and filled with internal solution for current clamp (130 mM K-Gluconate, 5 mM KCl, 10 mM HEPES, 2.5 mM MgCl2. 4 mM Na2ATP, 0.4 Na3GTP, 10 mM Na-phosphocreatine, 0.6 mM EGTA). After a giga-ohm seal was achieved, the membrane was disrupted with short bursts of negative pressure to achieve a whole-cell configuration. Rheobase current, the minimal current to elicit an action potential, was determined via stepwise injection of current in 25 pA increments. In optogenetic recordings, 470 nm LED pulses of 5 ms were delivered to activate ChR2 expressing neurons and to measure LED-induced postsynaptic potentials. The minimal LED % illumination required to generate consistent action potential firing varied between 5% - 15%. Recordings were made using a Scientifica SliceScope Pro 1000 and MultiClamp 700B (Molecular Device) with data filtered at 2.4 kHz and digitized at 10 kHz using a Digidata 1440A interface (Molecular Device) driven by pClamp 9.2 (Molecular Devices).

### Immunofluorescence

Mice were deeply anesthetized with 3% inhaled isoflurane (Kent Scientific; SomnoFlo) and perfused transcardially with 10 mL chilled phosphate buffered saline (PBS) followed by 10 mL chilled 4% paraformaldehyde (PFA) in PBS. Brains were then dissected and postfixed in 4% PFA in PBS at 4°C for 24 hours. Brains were then submerged in 30% sucrose for at least 24 hours prior to sectioning. Brains were sectioned using a Leica SM2010R Sliding Microtome at 60 μm. Sections were stored in cryoprotectant solution at -20°C, rinsed thoroughly in PBS and blocked in 5% donkey serum diluted in PBST (1X PBS with 0.3% TritonX-100) for 30 minutes at room temperature. Sections were then incubated in the primary antibody, diluted in PBST at 4°C: Guinea pig anti-NeuN (Millipore ABN90P, 1:1000) and Rabbit anti-HA (Cell Signaling, C29F4, 1:4000). ChR2-mCherry was visualized endogenously without antibody staining. Sections were rinsed with 1X PBS, mounted onto glass slides, dried, and cover slipped with Prolong Gold Antifade. Slides were imaged on a Keyence BZ-X710 fluorescence microscope. Further details about antibodies used in this study can be found in Supplemental Table S1.

### Ex vivo Optogenetic Activation

Acute brain slices were optogenetically activated using a custom built LED illumination system. This system consisted of a Ultra High Power LED (Prizmatix UHP-T-455-MP) with special 1-inch collimating optics for Microplate Illumination that could be remotely controlled with the benchtop UHP-T-LED Current Controller (Prizmatix) and the Pulser Plus (Prizmatix) to create trains of TTL pulses. The LED height was optimized for homogeneous light distribution to each well. All optogenetic stimulation described in the study consisted of 5 trains of blue light pulses (450–465 nm, peak at 455 nm) at 5 Hz for 30 seconds. Slices were allowed to recover in carbogenated aCSF for 30 minutes after optogenetic stimulation. For Opto-CLIP, slices were then crosslinked on ice, the CA1 hippocampus was rapidly dissected on ice, and tissue was processed for CLIP. For Opto-RiboTag, slices were transferred to cold HBSS + 0.1 mg/mL cycloheximide, the CA1 hippocampus was rapidly dissected on ice, and tissue was processed for RiboTag.

### FMRP-cTag CLIP

Each independent biological replicate (n=4) consisted of hippocampi pooled from 5 *Camk2a*-Cre;*Fmr1*-cTag mice aged 11 weeks. Crosslinked tissue was resuspended in 1.5 mL lysis buffer (1X PBS, 1% NP-40, 0.5% NaDOC and 0.1% SDS with cOmplete protease inhibitors (Roche)). Material was homogenized by mechanical homogenization and frozen at 80°C to ensure full cell lysis. Lysates were thawed and subject to DNase treatment (67.5 μl of RQ1 DNase (Promega) per 1.5 mL; 5 minutes; 37°C; 1100 rpm in thermomixer) and RNase treatment (final dilution of 1:1,666,666 of RNase A (Affymetrix); 5 minutes; 37°C; 1100 rpm in thermomixer). Lysate was then clarified by centrifugation at 20,000 x g for 20 min. The resulting supernatant was pre-cleared by rotating with 50 μl Protein G Dynabeads (Invitrogen) (washed in lysis buffer) for 45 min at 4°C. The pre-cleared supernatant was used for immunoprecipitation with 200 μl of Protein G Dynabeads (Invitrogen) loaded with 25 μg each of mouse monoclonal anti-GFP antibodies 19F7 and 19C8 (Heiman et al. 2008). Immunoprecipitation was performed for 2 hr at 4°C with rotation. Beads were then washed twice with lysis buffer, twice with high salt lysis buffer (5X PBS, 1% NP-40, 0.5% NaDOC and 0.1% SDS), twice with stringent wash buffer (15 mM Tris pH 7.5, 5 mM EDTA, 2.5 mM EGTA, 1% TritonX-100, 1% NaDOC, 0.1% SDS, 120 mM NaCl, 25 mM KCl), twice with high salt wash buffer (15 mM Tris pH 7.5, 5 mM EDTA, 2.5 mM EGTA, 1% TritonX-100, 1% NaDOC, 0.1% SDS, 1M NaCl), twice with low salt wash buffer (15 mM Tris pH 7.5, 5 mM EDTA), and twice with PNK wash buffer (50 mM Tris pH 7.4, 10 mM MgCl2, 0.5% NP-40). RNA tags were dephosphorylated with Alkaline Phosphatase (Roche) and subjected to overnight 3’ ligation at 16°C with a pre-adenylated linker (preA-L32) with the following ligation reaction: 2 μl of 25 mM linker, 2 μl of Truncated T4 RNA Ligase (NEB), 1X ligation buffer (NEB), 2 μl RNase inhibitor (Invitrogen), and 8 μl PEG8000 (NEB). The beads were washed three times with PNK wash buffer and the RNA-protein complexes labeled with ^32^P with the following reaction: 4 μl 10X PNK buffer (NEB), 2 μl T4 PNK enzyme, 1 μl ^32^P-γ-ATP at 37°C for 20 minutes shaking at 1100 rpm in a Thermomixer. Beads were then washed three times in PNK/ EGTA wash buffer (50 mM Tris pH 7.4, 20 mM EGTA, 0.5% NP-40) and incubated at 70°C for 10 minutes shaking at 1100 rpm in a Thermomixer to eluate RNA/protein complexes from beads and subjected to SDS-PAGE and transfer as described (Moore et al. 2014). Regions corresponding to 120-175 kDa were excised from the nitrocellulose membrane and RNA tags were collected with proteinase K/Urea followed by chloroform:isoamyl alcohol (24:1) treatment as previously described (Hwang et al. 2017). Cloning was performed using the BrdU-CLIP protocol as described (Sawicka et al. 2019; Hale et al. 2021) using RT primers with six nucleotide barcode index sequences to allow for up to 24 samples to be pooled together in one MiSeq sequencing run (Illumina) and subsequently demultiplexed. CLIP reads were processed as described previously using CLIP Tool Kit (CTK) (Moore et al. 2014; Shah et al. 2017). Briefly, raw reads were filtered for quality and demultiplexed using indexes introduced during the reverse transcription reaction. PCR duplicates were collapsed and adapter sequences removed. Reads were mapped to the mm10 RefSeq genome using BWA (Dobin et al. 2013). Mapped reads were further collapsed using fastq2collapse.pl from the CTK toolkit with default options. The distribution of CLIP reads that uniquely map to the genome (Fig. 4G) was determined using bed2annotation.pl from the CTK toolkit. Statistical correlation between samples was assessed using the Pearson’s product moment correlation coefficient (cor.test in R).

### TRAP- and RNA-seq of acute brain slices

Each independent biological replicate (n=3) consisted of hippocampi pooled from *Camk2a*-Cre; *Rpl22*-HA mice aged 11 weeks. Dissected hippocampi from acute brain slices were pooled, resuspended in 5% w/v ice-cold polysome buffer (20 mM HEPES, pH 7.4, 150 mM NaCl, 5 mM MgCl2, 0.5 mM DTT) freshly supplemented with 40 U/mL RNasin Plus (Promega), cOmplete protease inhibitors (Roche), 0.1 mg/mL cycloheximide, and 0.5 mM DTT) and homogenized by dounce homogenization (40 strokes type A pestle followed by 40 strokes type B pestle). NP-40 was added to 1% final concentration and lysate was incubated on ice for 10 min. Supernatant was subsequently centrifuged at 2000 x g for 10 min at 4°C followed by 20,000 x g for 10 min at 4°C. 10% of the resulting lysate was used as input and the remaining lysate was pre-cleared by rotation with 50 μl Protein G Dynabeads (Invitrogen) (washed in polysome buffer) for 45 min at 4°C. HA-tagged ribosomes were collected by indirect IP by adding mouse anti-HA (8 μl antibody per mL lysate; Biolegend; MMS-101R) antibody directly to the lysate, rotating the mixture at 4°C for 4 hours, and adding 150 μl Protein G dynabeads to the lysate + antibody followed by overnight rotation at 4°C. Beads were washed twice with polysome buffer containing 1% NP-40 and three times with high salt wash buffer (20 mM HEPES, 5 mM MgCl2, 350 mM KCl, 1% NP-40) freshly supplemented with 0.1 mg/mL cycloheximide, and 1 mM DTT. All washes were 5 minutes with rotation at 4°C. RNA was extracted from input and IP samples by incubation with 1 mL Trizol at room temperature for 5 minutes and then RNeasy Lipid Tissue Mini Kit (Qiagen) via the manufacturer’s protocol with on-column DNase treatment (Roche). RNA quantity and quality were obtained by the Agilent 2100 Pico Bioanalyzer system. The libraries were prepared by Illumina Stranded Total RNA Prep with Ribo-Zero Plus following the manufacturer’s instructions. High-throughput sequencing was performed on NovaSeq (Illumina) to obtain 150 nucleotide paired-end reads. Transcript expression was quantified Input and IP samples using salmon (Patro et al. 2017) and mm10 gene models. Differential expression analysis was performed using DESeq2 (Love et al. 2014). Transcripts were considered significantly enriched (IP vs input; activated vs control) with a Benjamini–Hochberg p.adjusted less than 0.05.

### CLIP normalization using RiboTag

RiboTag transcript per million (TPM) was determined for each transcript using salmon (Patro et al. 2017) and a single transcript with the highest mean TPM across all activated and control replicates was selected per gene. This set of 14353 transcripts was used for all subsequent analyses. CLIP tags were preprocessed as described above and then were mapped to the transcriptome using STAR (Dobin et al. 2013) with --quantMode GeneCounts TranscriptomeSAM. Mapped reads were collapsed using fastq2collapse.pl from the CTK toolkit with default options. Only reads that mapped to transcripts expressed by RiboTag were retained. CLIP scores were calculated as described previously (Sawicka et al. 2019; Hale et al. 2021). Briefly, for each replicate, CLIP TPM was calculated based on transcript length and library size. Scatter plots of log_2_ CLIP TPM vs log_2_ RiboTag TPM for each condition and replicate were made and a linear regression line fitted to the data. The CLIP score for each transcript and each replicate was then calculated from the slope (lm slope) and intercept (lm intercept) of the fitted line using the equation: CLIP score = log_2_ CLIP TPM -(lm slope x RiboTag TPM) + lm intercept. The final CLIP score for each transcript was determined as the mean CLIP score across all biological replicates. Transcripts were classed as stringent targets if they had a CLIP score greater than 2 across all biological replicates, high binding targets if they had a CLIP score greater than 1 but less than 2, low binding targets if they had a CLIP score between 0 and 1. All other transcripts were classified as non-targets. Differential comparison of CLIP scores between control and activated conditions was done with limma (Ritchie et al. 2015).

### Gene Ontology (GO) and Gene Set Enrichment Analysis (GSEA)

Gene ontology (GO) analysis was performed using the function enrichGO in the R package clusterProfiler (Yu et al. 2012). GSEA was done using the function gseGO in the R package clusterProfiler (Yu et al. 2012). The input for GSEA was log_2_ fold change of CLIP scores in activated versus control conditions determined by limma (Ritchie et al. 2015).

## Data and Code availability

Sequencing data for FMRP-CLIP have been deposited in GEO under accession code GSE286379. Sequencing data for Opto-RiboTag have been deposited in GEO under accession code GSE286381. Further details about software used in this study can be found in Supplemental Table S1. All custom R scripts used in this study have been deposited on the public Github repository: https://github.com/ruthasinger

## ACKNOWLEDGMENTS

We thank C. Zhao, C. Lai, and B. Zhang from the Genomics Core and T. Carroll, D. Barrows, and W. Wang from the Bioinformatics Core for their insights and technical support. We thank members of the Darnell laboratory at The Rockefeller University, particularly E. Hacisuleyman, for thoughtful discussions and review of the paper. This work was supported by Glenn AFAR Foundation and NINDS F32NS119376 Postdoctoral Fellowships (R.A.S) and NINDS R35NS097404 (R.B.D). R.B.D. is an Investigator of the Howard Hughes Medical Institute.

## AUTHOR CONTRIBUTIONS

Conceptualization, R.A.S. and R.B.D.; Data Curation, R.A.S.; Formal Analysis, R.A.S.; Funding Acquisition, R.A.S. and R.B.D.; Investigation, R.A.S., V.R., and K.P.; Methodology, R.A.S. and R.B.D.; Supervision, N.H. and R.B.D.; Visualization, R.A.S. and R.B.D.; Writing – Original Draft Preparation, R.A.S. and R.B.D.; Writing – Review & Editing, R.A.S., V.R., K.P., N.H., and R.B.D.

---

**Supplemental Figure 1.**
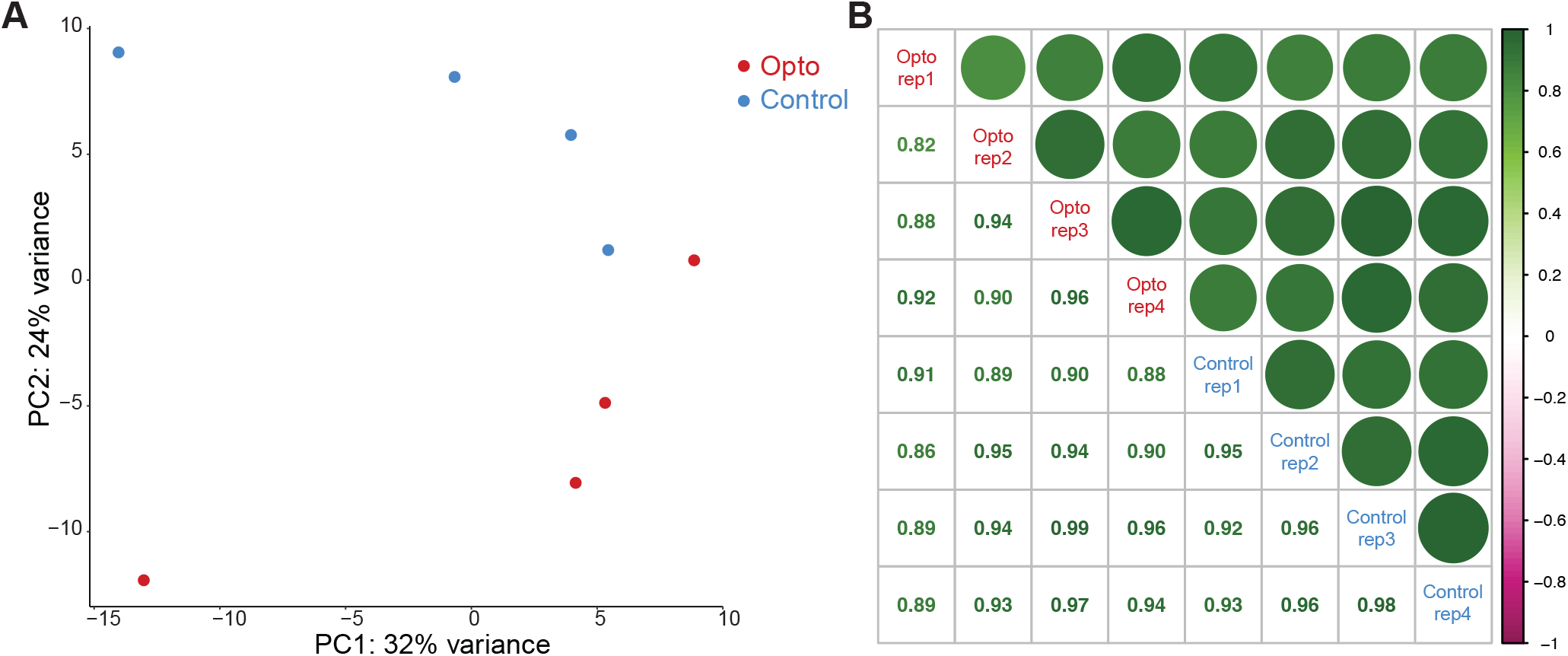
A) Principal component analysis (PCA) shown for Opto-FMRP-CLIP samples (red) and Control FMRP-CLIP samples (blue). B) Correlation plots of each CLIP sample compared to each other. Color of the circle corresponds to the r-squared value, which is listed in the matching box on the other half of the matrix. Opto samples are listed in red text and Control samples are in blue text (n=4).

**Supplemental Figure 2.**
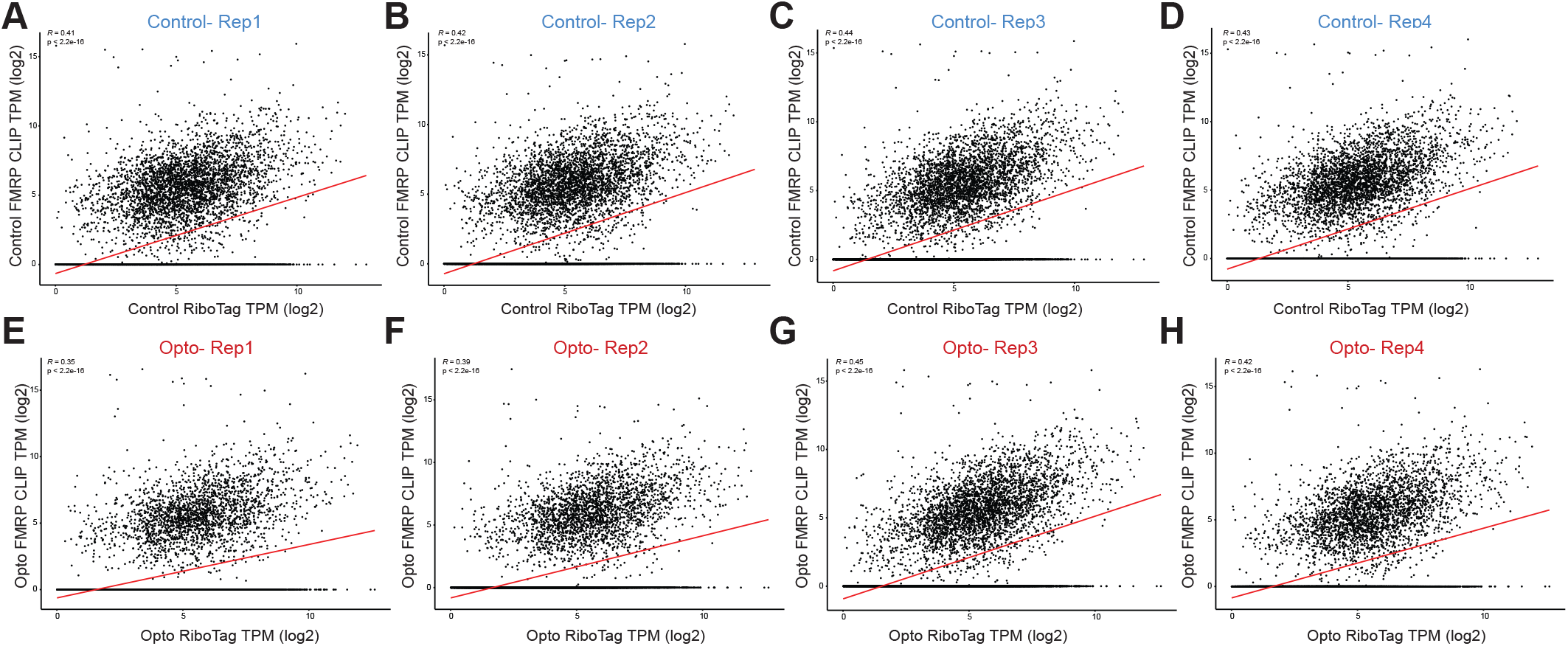
A and B) FMRP-CLIP targets were defined by normalizing CLIP tag density across the coding region to the relative abundance of the transcript as measured by RiboTag. A CLIP score per transcript was calculated independently for each of the eight CLIP biological replicates. Scatter plots of log_2_ normalized transcript per million (TPM) of CLIP tags from an individual Control (A) or Opto-FMRP-CLIP (B) experiment compared to average log_2_ RiboTag TPM from Control (A) or Opto-RiboTag (B). n=4 biological replicates for FMRP-CLIP. n=3 biological replicates for RiboTag. Red line indicates the linear regression model used to calculate CLIP scores. See methods for further details on CLIP score calculations.

**Supplemental Figure 3.**
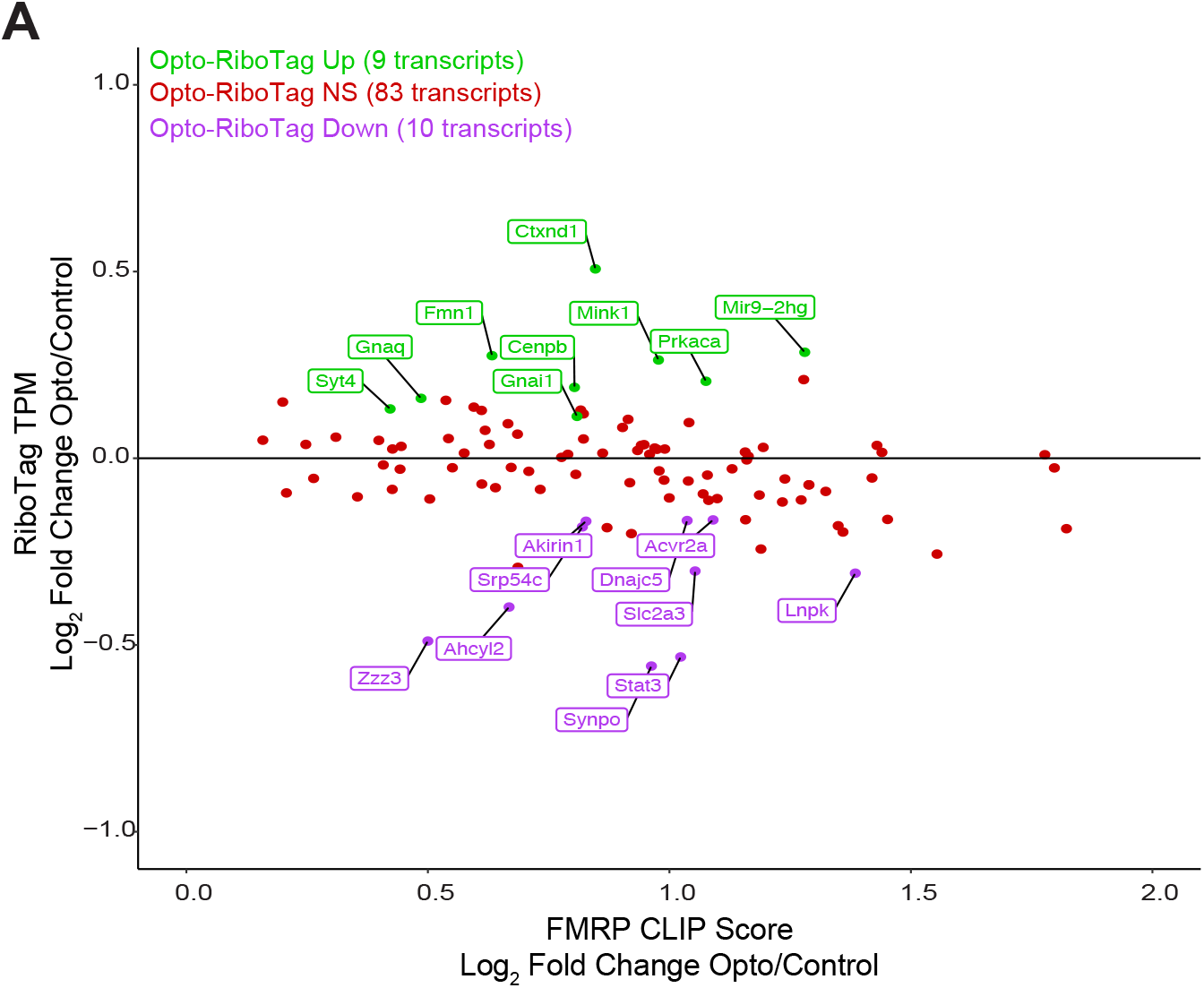
A) Scatter plot of Opto-RiboTag versus Control RiboTag (log_2_ TPM) compared to Op-to-CLIP versus Control CLIP CLIP scores (log_2_ fold change). Red dots indicate transcripts that are more bound by FMRP in activated neurons (FMRP-Up) and are unchanged by Opto RiboTag. Green dots indicate transcripts that are more bound by FMRP in activated neurons (FMRP-Up) and are upregulated in activated RiboTag. Purple dots indicate transcripts that are more bound by FMRP in activated neurons (FMRP-Up) and are downregulated in activated RiboTag.

**Supplemental Table 1.**
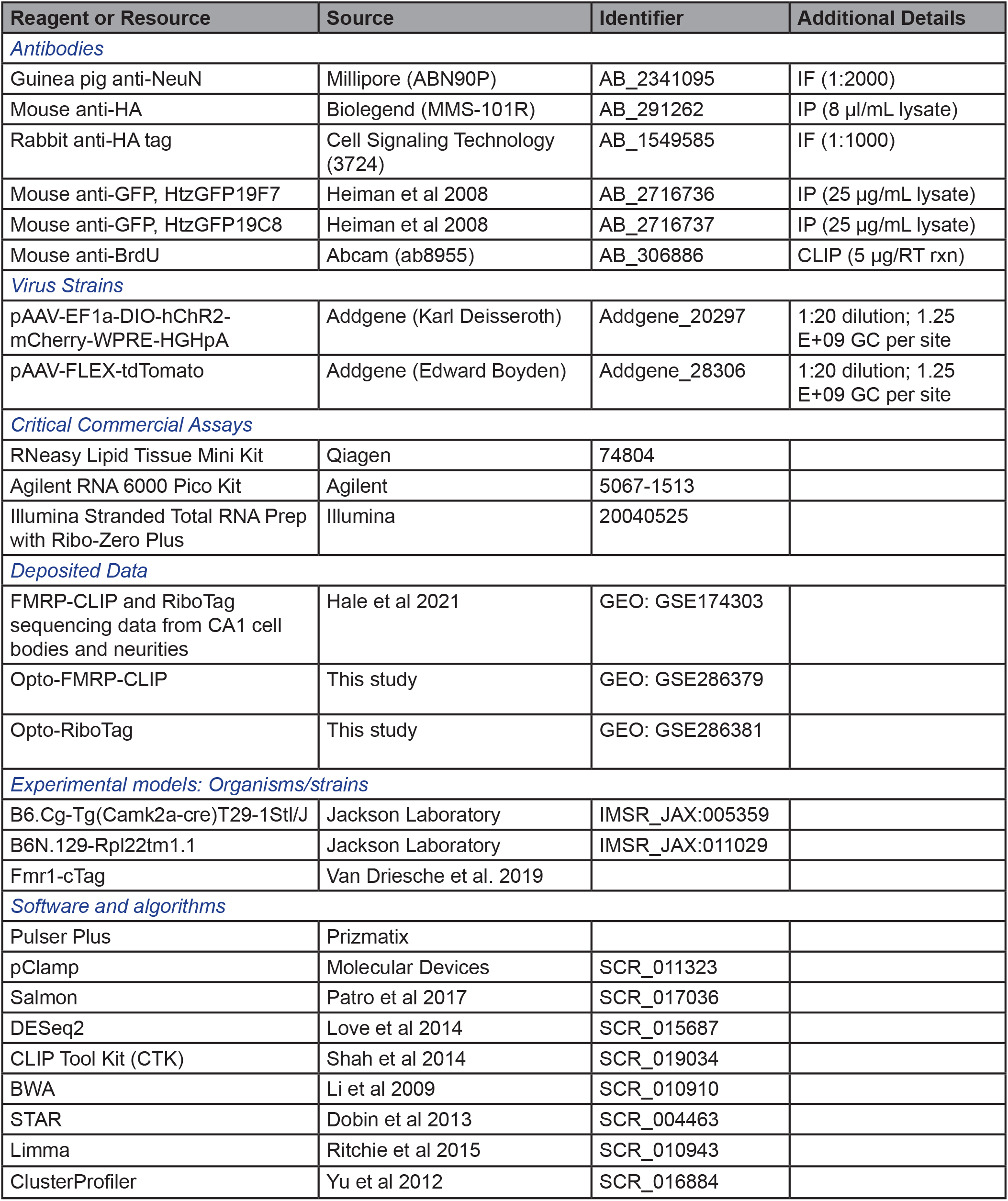
Key resources used in this study.

## Notes

### Competing Interest Statement

The authors have declared no competing interest.

### Summary of Updates

We have added a new Figure 7 and Supplemental Figure S3. Some of the text has been updated to reflect new analyses done.

## REFERENCES

Anadolu MN, Sun J, Kailasam S, Chalkiadaki K, Krimbacher K, Li JT-Y, Markova T, Jafarnejad SM, Lefebvre F, Ortega J, et al. 2023. Ribosomes in RNA Granules Are Stalled on mRNA Sequences That Are Consensus Sites for FMRP Association. J Neurosci 43: 2440–2459.

Arbab T, Pennartz CMA, Battaglia FP. 2018. Impaired hippocampal representation of place in the Fmr1-knockout mouse model of fragile X syndrome. Sci Rep 8: 8889.

Aryal S, Klann E. 2018. Turning up translation in fragile X syndrome. Science 361: 648–649.

Ashley CT Jr, Wilkinson KD, Reines D, Warren ST. 1993. FMR1 protein: conserved RNP family domains and selective RNA binding. Science 262: 563–566.

Bear MF, Huber KM, Warren ST. 2004. The mGluR theory of fragile X mental retardation. Trends Neurosci 27: 370–377.

Benito E, Barco A. 2015. The neuronal activity-driven transcriptome. Mol Neurobiol 51: 1071–1088.

Boone CE, Davoudi H, Harrold JB, Foster DJ. 2018. Abnormal Sleep Architecture and Hippocampal Circuit Dysfunction in a Mouse Model of Fragile X Syndrome. Neuroscience 384: 275–289.

Cavallaro S, D’Agata V, Manickam P, Dufour F, Alkon DL. 2002. Memory-specific temporal profiles of gene expression in the hippocampus. Proc Natl Acad Sci U S A 99: 16279–16284.

Chen PB, Kawaguchi R, Blum C, Achiro JM, Coppola G, O’Dell TJ, Martin KC. 2017. Mapping Gene Expression in Excitatory Neurons during Hippocampal Late-Phase Long-Term Potentiation. Front Mol Neurosci 10: 39.

Comery TA, Harris JB, Willems PJ, Oostra BA, Irwin SA, Weiler IJ, Greenough WT. 1997. Abnormal dendritic spines in fragile X knockout mice: Maturation and pruning deficits. Proceedings of the National Academy of Sciences 94: 5401–5404.

Darnell JC, Van Driesche SJ, Zhang C, Hung KYS, Mele A, Fraser CE, Stone EF, Chen C, Fak JJ, Chi SW, et al. 2011. FMRP stalls ribosomal translocation on mRNAs linked to synaptic function and autism. Cell 146: 247–261.

Darnell RB. 2020. The Genetic Control of Stoichiometry Underlying Autism. Annu Rev Neurosci 43: 509–533.

Deisseroth K, Feng G, Majewska AK, Miesenböck G, Ting A, Schnitzer MJ. 2006. Next-Generation Optical Technologies for Illuminating Genetically Targeted Brain Circuits. J Neurosci 26: 10380–10386.

Dobin A, Davis CA, Schlesinger F, Drenkow J, Zaleski C, Jha S, Batut P, Chaisson M, Gingeras TR. 2013. STAR: ultrafast universal RNA-seq aligner. Bioinformatics 29: 15–21.

Doyle JP, Dougherty JD, Heiman M, Schmidt EF, Stevens TR, Ma G, Bupp S, Shrestha P, Shah RD, Doughty ML, et al. 2008. Application of a translational profiling approach for the comparative analysis of CNS cell types. Cell 135: 749–762.

French PJ, O’Connor V, Voss K, Stean T, Hunt SP, Bliss TV. 2001. Seizure-induced gene expression in area CA1 of the mouse hippocampus. Eur J Neurosci 14: 2037–2041.

Frey U, Krug M, Brödemann R, Reymann K, Matthies H. 1989. Long-term potentiation induced in dendrites separated from rat’s CA1 pyramidal somata does not establish a late phase. Neurosci Lett 97: 135–139.

Guzowski JF, McNaughton BL, Barnes CA, Worley PF. 1999. Environment-specific expression of the immediate-early gene Arc in hippocampal neuronal ensembles. Nat Neurosci 2: 1120– 1124.

Hacisuleyman E, Hale CR, Noble N, Luo J-D, Fak JJ, Saito M, Chen J, Weissman JS, Darnell RB. 2024. Neuronal activity rapidly reprograms dendritic translation via eIF4G2:uORF binding. Nat Neurosci 27: 822–835.

Hagerman RJ, Berry-Kravis E, Hazlett HC, Bailey DB Jr, Moine H, Kooy RF, Tassone F, Gantois I, Sonenberg N, Mandel JL, et al. 2017. Fragile X syndrome. Nat Rev Dis Primers 3: 17065.

Hale CR, Sawicka K, Mora K, Fak JJ, Kang JJ, Cutrim P, Cialowicz K, Carroll TS, Darnell RB. 2021. FMRP regulates mRNAs encoding distinct functions in the cell body and dendrites of CA1 pyramidal neurons. Elife 10:e71892.

Heiman M, Kulicke R, Fenster RJ, Greengard P, Heintz N. 2014. Cell type-specific mRNA purification by translating ribosome affinity purification (TRAP). Nat Protoc 9: 1282–1291.

Heiman M, Schaefer A, Gong S, Peterson JD, Day M, Ramsey KE, Suárez-Fariñas M, Schwarz C, Stephan DA, Surmeier DJ, et al. 2008. A translational profiling approach for the molecular characterization of CNS cell types. Cell 135: 738–748.

Huber KM, Gallagher SM, Warren ST, Bear MF. 2002. Altered synaptic plasticity in a mouse model of fragile X mental retardation. Proceedings of the National Academy of Sciences 99: 7746–7750.

Huber KM, Kayser MS, Bear MF. 2000. Role for rapid dendritic protein synthesis in hippocampal mGluR-dependent long-term depression. Science 288: 1254–1257.

Hughes JR. 1958. Post-tetanic potentiation. Physiol Rev 38: 91–113.

Hwang H-W, Saito Y, Park CY, Blachère NE, Tajima Y, Fak JJ, Zucker-Scharff I, Darnell RB. 2017. cTag-PAPERCLIP Reveals Alternative Polyadenylation Promotes Cell-Type Specific Protein Diversity and Shifts Araf Isoforms with Microglia Activation. Neuron 95: 1334–1349.

Jaeger BN, Linker SB, Parylak SL, Barron JJ, Gallina IS, Saavedra CD, Fitzpatrick C, Lim CK, Schafer ST, Lacar B, et al. 2018. A novel environment-evoked transcriptional signature predicts reactivity in single dentate granule neurons. Nat Commun 9: 3084.

Jereb S, Hwang H-W, Van Otterloo E, Govek E-E, Fak JJ, Yuan Y, Hatten ME, Darnell RB. 2018. Differential 3’ processing of specific transcripts expands regulatory and protein diversity across neuronal cell types. Elife 7: e34042.

Kang H, Schuman EM. 1996. A requirement for local protein synthesis in neurotrophin-induced hippocampal synaptic plasticity. Science 273: 1402–1406.

Kazdoba TM, Leach PT, Silverman JL, Crawley JN. 2014. Modeling fragile X syndrome in the Fmr1 knockout mouse. Intractable Rare Dis Res 3: 118–133.

Kim T-K, Hemberg M, Gray JM, Costa AM, Bear DM, Wu J, Harmin DA, Laptewicz M, Barbara-Haley K, Kuersten S, et al. 2010. Widespread transcription at neuronal activity-regulated enhancers. Nature 465: 182–187.

Korb E, Herre M, Zucker-Scharff I, Gresack J, Allis CD, Darnell RB. 2017. Excess Translation of Epigenetic Regulators Contributes to Fragile X Syndrome and Is Alleviated by Brd4 Inhibition. Cell 170: 1209–1223.

Kuwajima M, Ostrovskaya OI, Cao G, Weisberg SA, Harris KM, Zemelman BV. 2020. Ultra-structure of light-activated axons following optogenetic stimulation to produce late-phase long-term potentiation. PLoS One 15: e0226797.

Lauterborn JC, Rex CS, Kramár E, Chen LY, Pandyarajan V, Lynch G, Gall CM. 2007. Brain-derived neurotrophic factor rescues synaptic plasticity in a mouse model of fragile X syndrome. J Neurosci 27: 10685–10694.

Lee HY, Ge W-P, Huang W, He Y, Wang GX, Rowson-Baldwin A, Smith SJ, Jan YN, Jan LY. 2011. Bidirectional regulation of dendritic voltage-gated potassium channels by the fragile X mental retardation protein. Neuron 72: 630–642.

Lin JY. 2011. A user’s guide to channelrhodopsin variants: features, limitations and future developments: A user’s guide to channelrhodopsin variants. Exp Physiol 96: 19–25.

Love MI, Huber W, Anders S. 2014. Moderated estimation of fold change and dispersion for RNA-seq data with DESeq2. Genome Biol 15: 550.

Moore MJ, Blachere NE, Fak JJ, Park CY, Sawicka K, Parveen S, Zucker-Scharff I, Moltedo B, Rudensky AY, Darnell RB. 2018. ZFP36 RNA-binding proteins restrain T cell activation and anti-viral immunity. Elife 7:e33057.

Moore MJ, Zhang C, Gantman EC, Mele A, Darnell JC, Darnell RB. 2014. Mapping Argonaute and conventional RNA-binding protein interactions with RNA at single-nucleotide resolution using HITS-CLIP and CIMS analysis. Nat Protoc 9: 263–293.

Muddashetty RS, Kelić S, Gross C, Xu M, Bassell GJ. 2007. Dysregulated metabotropic glutamate receptor-dependent translation of AMPA receptor and postsynaptic density-95 mRNAs at synapses in a mouse model of fragile X syndrome. J Neurosci 27: 5338–5348.

Nalavadi VC, Muddashetty RS, Gross C, Bassell GJ. 2012. Dephosphorylation-induced ubiquitination and degradation of FMRP in dendrites: a role in immediate early mGluR-stimulated translation. J Neurosci 32: 2582–2587.

Narayanan U, Nalavadi V, Nakamoto M, Pallas DC, Ceman S, Bassell GJ, Warren ST. 2007. FMRP phosphorylation reveals an immediate-early signaling pathway triggered by group I mGluR and mediated by PP2A. J Neurosci 27: 14349–14357.

Park CS, Gong R, Stuart J, Tang S-J. 2006. Molecular network and chromosomal clustering of genes involved in synaptic plasticity in the hippocampus. J Biol Chem 281: 30195–30211.

Patro R, Duggal G, Love MI, Irizarry RA, Kingsford C. 2017. Salmon provides fast and bias-aware quantification of transcript expression. Nat Methods 14: 417–419.

Ritchie ME, Phipson B, Wu D, Hu Y, Law CW, Shi W, Smyth GK. 2015. limma powers differential expression analyses for RNA-sequencing and microarray studies. Nucleic Acids Res 43: e47.

Saito Y, Yuan Y, Zucker-Scharff I, Fak JJ, Jereb S, Tajima Y, Licatalosi DD, Darnell RB. 2019. Differential NOVA2-Mediated Splicing in Excitatory and Inhibitory Neurons Regulates Cortical Development and Cerebellar Function. Neuron 101: 707–720.

Sanz E, Yang L, Su T, Morris DR, McKnight GS, Amieux PS. 2009. Cell-type-specific isolation of ribosome-associated mRNA from complex tissues. Proc Natl Acad Sci U S A 106: 13939– 13944.

Sawicka K, Hale CR, Park CY, Fak JJ, Gresack JE, Van Driesche SJ, Kang JJ, Darnell JC, Darnell RB. 2019. FMRP has a cell-type-specific role in CA1 pyramidal neurons to regulate autism-related transcripts and circadian memory. Elife 8:e46919.

Scanziani M, Häusser M. 2009. Electrophysiology in the age of light. Nature 461: 930–939.

Shah A, Qian Y, Weyn-Vanhentenryck SM, Zhang C. 2017. CLIP Tool Kit (CTK): a flexible and robust pipeline to analyze CLIP sequencing data. Bioinformatics 33: 566–567.

Talbot ZN, Sparks FT, Dvorak D, Curran BM, Alarcon JM, Fenton AA. 2018. Normal CA1 Place Fields but Discoordinated Network Discharge in a Fmr1-Null Mouse Model of Fragile X Syndrome. Neuron 97: 684–697.

Thomson SR, Seo SS, Barnes SA, Louros SR, Muscas M, Dando O, Kirby C, Wyllie DJA, Hardingham GE, Kind PC, et al. 2017. Cell-Type-Specific Translation Profiling Reveals a Novel Strategy for Treating Fragile X Syndrome. Neuron 95: 550–563.

Tsien JZ, Chen DF, Gerber D, Tom C, Mercer EH, Anderson DJ, Mayford M, Kandel ER, Tonegawa S. 1996. Subregion- and Cell Type–Restricted Gene Knockout in Mouse Brain. Cell 87: 1317–1326.

Turrigiano GG, Nelson SB. 2004. Homeostatic plasticity in the developing nervous system. Nat Rev Neurosci 5: 97–107.

Van Driesche SJ, Sawicka K, Zhang C, Hung SKY, Park CY, Fak JJ, Yang C, Darnell RB, Darnell JC. 2019. FMRP binding to a ranked subset of long genes is revealed by coupled CLIP and TRAP in specific neuronal cell types. bioRxiv doi: 10.1101/762500

Weyn-Vanhentenryck SM, Mele A, Yan Q, Sun S, Farny N, Zhang Z, Xue C, Herre M, Silver PA, Zhang MQ, et al. 2014. HITS-CLIP and integrative modeling define the Rbfox splicing-regulatory network linked to brain development and autism. Cell Rep 6: 1139–1152.

Yizhar O, Fenno LE, Davidson TJ, Mogri M, Deisseroth K. 2011. Optogenetics in neural systems. Neuron 71: 9–34.

Yu G, Wang L-G, Han Y, He Q-Y. 2012. clusterProfiler: an R package for comparing biological themes among gene clusters. OMICS 16: 284–287.

